# CD8^+^ T cells are primed by cDC1 and exacerbate tau-mediated neurodegeneration

**DOI:** 10.64898/2026.02.26.708260

**Authors:** Hao Hu, Peter Bor-Chian Lin, Carisa Zeng, Prabal Sharma, Yongyi Li, Hong Jiang, Jonathan Nulman, Ray A. Ohara, Tong Wu, Shasha Li, Wayne M. Yokoyama, Maxim N. Artyomov, Kenneth M. Murphy, Jason D. Ulrich, David M. Holtzman

## Abstract

There are changes in adaptive immunity in Alzheimer’s disease (AD) and increases in activated CD8^+^ T cells in brain correlate with tau pathology^1–3^. However, which cells mediate T cell priming in tau-mediated neurodegeneration remains unclear. In different conditions such as cancer, viral infections, and autoimmune diseases outside the CNS, conventional type-1 dendritic cells (cDC1) perform antigen cross-presentation to prime CD8^+^ T cells^4,5^. We demonstrate that tauopathy mice deficient in cDC1 are markedly protected against tau-mediated neurodegeneration and display a selective decrease in brain CD8^+^ T cell infiltration and glial reactivity. The remaining CD8^+^ T cells showed an antigen inexperienced status with less clonal expansion, indicating suboptimal T cell priming. We confirm that brain derived antigens are presented in secondary lymphoid tissues to prime CD8^+^ T cells. Our study identifies cDC1 cells as critical for CD8^+^ T cell priming outside the CNS. This priming is required for a large increase of activated CD8^+^ T cells in the brain which promotes tau-mediated neurodegeneration.

## Main text

Tauopathies are a group of neurodegenerative diseases, including Alzheimer’s disease (AD) and certain forms of frontal temporal dementia among others. They are defined by the intracellular fibrillar aggregation of hyperphosphorylated tau protein which tightly correlates with neuronal dysfunction and death^6^. Tauopathy, as well as other forms of AD pathologies, is accompanied by increases in reactive microglia and astrocytes as well as increases in T cells in the brain, particularly CD8^+^ cytotoxic T lymphocytes (CTLs)^1–3,7–10^. We previously found that depleting T cells significantly ameliorated tau-mediated neurodegeneration in P301S tau transgenic mice expressing human APOE4 (TE4), supporting an active role for T cells in driving tau-mediated neurodegeneration^1^. Both CD4^+^ and CD8^+^ T cells are increased in brain regions with tau pathology, but we were intrigued by the observation that clonally expanded CD8^+^ T cells constituted a majority of the infiltrating CD3^+^ T cells (∼60-70%) in the diseased brain. Conventional type 1 dendritic cells (cDC1) cells are specialized antigen presenting cells (APCs) that perform antigen cross-presentation, a process that involves presenting exogenously derived antigens on MHC-I molecules to prime CD8^+^ T cells^11–14^. In the context of brain specific T cell responses during neurodegeneration, it is not known whether cDC1 cells are responsible for cross-priming brain specific CD8^+^ T cells, ultimately leading to T cell infiltration into the brain as well as subsequent neuronal dysfunction and loss. Here we evaluate the role of cDC1 by genetically ablating these cells in TE4 mice. We found that eliminating cDC1 strongly protects against tau-mediated neurodegeneration and selectively affects CD8^+^ T cell accumulation in the diseased brain. Importantly, we also show that cDC1-dependent antigen cross-presentation of a brain-derived antigen occurs outside the CNS tissues, suggesting a therapeutic strategy by targeting this aspect of the peripheral immune system to treat tauopathies including AD.

### Knockout of *Irf8* +32-kb enhancer specifically eliminates cDC1 throughout the development of tau-mediated neurodegeneration

cDC1 development depends upon a cDC1-specific super-enhancer +32-kb of the *Irf8* gene^15^. cDC1 cells were shown to be stringently absent in *Irf8*+32^-/-^ mice even under pro-inflammatory conditions, without affecting Irf8 function in other leukocytes^15,16^. We crossed TE4 mice with *Irf8*+32^-/-^ to generate cDC1 deficient mice under the background of TE4 (TE4^Δ+32^). Four cohorts of mice were generated for further analysis: E4^WT^ (non-tau, cDC1 sufficient), E4^Δ+32^ (non-tau, cDC1 deficient), TE4^WT^ (tau expressing, cDC1 sufficient), and TE4^Δ+32^ (tau expressing, cDC1 deficient) (Extended Data Fig. 1a). We first assessed cDC1 development in 9.5-month-old TE4 mice, which exhibit robust tauopathy, reactive gliosis, and neurodegeneration, by sorting common dendritic cell progenitors (CDP), specified pre-cDC1 progenitors (pre-DC1), and pre-cDC2 progenitors (pre-DC2) from bone marrow and culturing these cells with Flt3L *ex vivo* for 7 days (Extended Data Fig. 1b,c). CDP derived from TE4^WT^ mice developed into both mature cDC1 and cDC2 cells and pre-DC1 only developed into cDC1 whereas pre-DC2 only developed into cDC2 (Extended Data Fig. 1d). As expected, CDP from TE4^Δ+32^ failed to generate cDC1. Even though CDP still specified to pre-DC1 in TE4^Δ+32^ bone marrow, pre-DC1 derived from TE4^Δ+32^ did not survive to further develop into cDC1. Development of cDC2 was normal in TE4^Δ+32^ mice (Extended Data Fig. 1d). As a result, cDC1 were not detectable in various tissues including spleen, deep cervical lymph nodes (dCLN) (the draining lymph nodes for brain), meninges, and brain parenchymal tissues in TE4^Δ+32^ mice (Extended Data Fig. 1e,f). cDC1 constituted a minor population (20-30%) of conventional DC cells and their percentage of total leukocytes was even smaller (accounting for ∼0.4-0.6% of leukocytes) (Extended Data Fig. 1g).

Irf8 is a critical transcription factor contributing microglia development and promoting the reactive states of microglia^17–20^. To exclude the possibility that deletion of +32-kb super enhancer also affected microglia responses during tau-mediated neurodegeneration, we compared the chromatin accessibility at the *Irf8* +32-kb enhancer in cDC1 and microglia as well as T cells from publicly available dataset (https://www.immgen.org/)^21^. As with previous findings^15,21^, the *Irf8*+32-kb enhancer was only accessible in cDC1 cells but not in microglia or T cells, while the macrophage-specific *Irf8*-50-kb enhancer was accessible in microglia, but not in cDC1 cells (Extended Data Fig. 2a)^21^. Consistent with this finding, we performed qRT-PCR analysis to detect *Irf8* expression at the RNA level in sorted microglia (Extended Data Fig. 2b) and did not observe a difference in *Irf8* expression in *Irf8*+32^-/-^ mice (both E4^Δ+32^ and TE4^Δ+32^) (Extended Data Fig. 2c). Furthermore, Irf8 expression at the protein level was also intact in E4^Δ+32^, comparable with E4^WT^ (Extended Data Fig. 2d).

### cDC1 deficient TE4 mice are protected against tau-mediated neurodegeneration

We next assessed the effect of cDC1 deficiency on neuropathology in 9.5-month-old TE4 mice. Male and female TE4^WT^ mice displayed severe regional brain atrophy with marked volume loss in both the hippocampus and piriform/entorhinal cortex (P-E) regions as well as enlargement of the lateral ventricles (LV) compared to E4^WT^ (Fig. 1a,b). cDC1 deficient mice showed significant preservation of the hippocampus and P-E tissues and a trending reduction in size of the LV (Fig. 1a,b) compared to TE4^WT^. Likewise, the dentate gyrus (DG) granule cell layer was significantly thinner in male and female TE4^WT^ compared to E4^WT^ mice. This thinning was rescued in TE4^Δ+32^ mice (Fig.1c,d). Plasma neurofilament light chain (NfL) level is a reliable biomarker of neurodegeneration^1^. Plasma NfL levels were significantly lower in male TE4^Δ+32^ than in TE4^WT^ mice but were not significantly different in female TE4^Δ+32^ compared to TE4^WT^ mice (Fig.1e). Nest building is an innate behavior that is negatively impacted by tau-dependent neurodegeneration in P301S mice^22,23^. Both male and female TE4^Δ+32^ mice had significantly higher nest-building scores than TE4^WT^ mice (Extended Data Fig. 3a,b), suggesting better cognitive function. Consistent with brain volume data, the overall brain weight was reduced in male and female TE4^WT^ mice compared to E4^WT^, but TE4^Δ+32^ mice had significantly increased brain weights compared to TE4^WT^ mice (Extended Data Fig. 3c). Brain weights positively correlated with hippocampal volume (Extended Data Fig. 3d). Surprisingly, the level of cDC1 in brain parenchymal tissues remained extremely low, and cDC1 were barely detectable in the brains of 9.5-month-old TE4^WT^ mice even when brain atrophy was readily apparent (Extended Data Fig. 1f,g). These results indicated that despite their paucity in the brain parenchyma, cDC1 cells played an active role in promoting tau-mediated neurodegeneration.

**Fig. 1.**
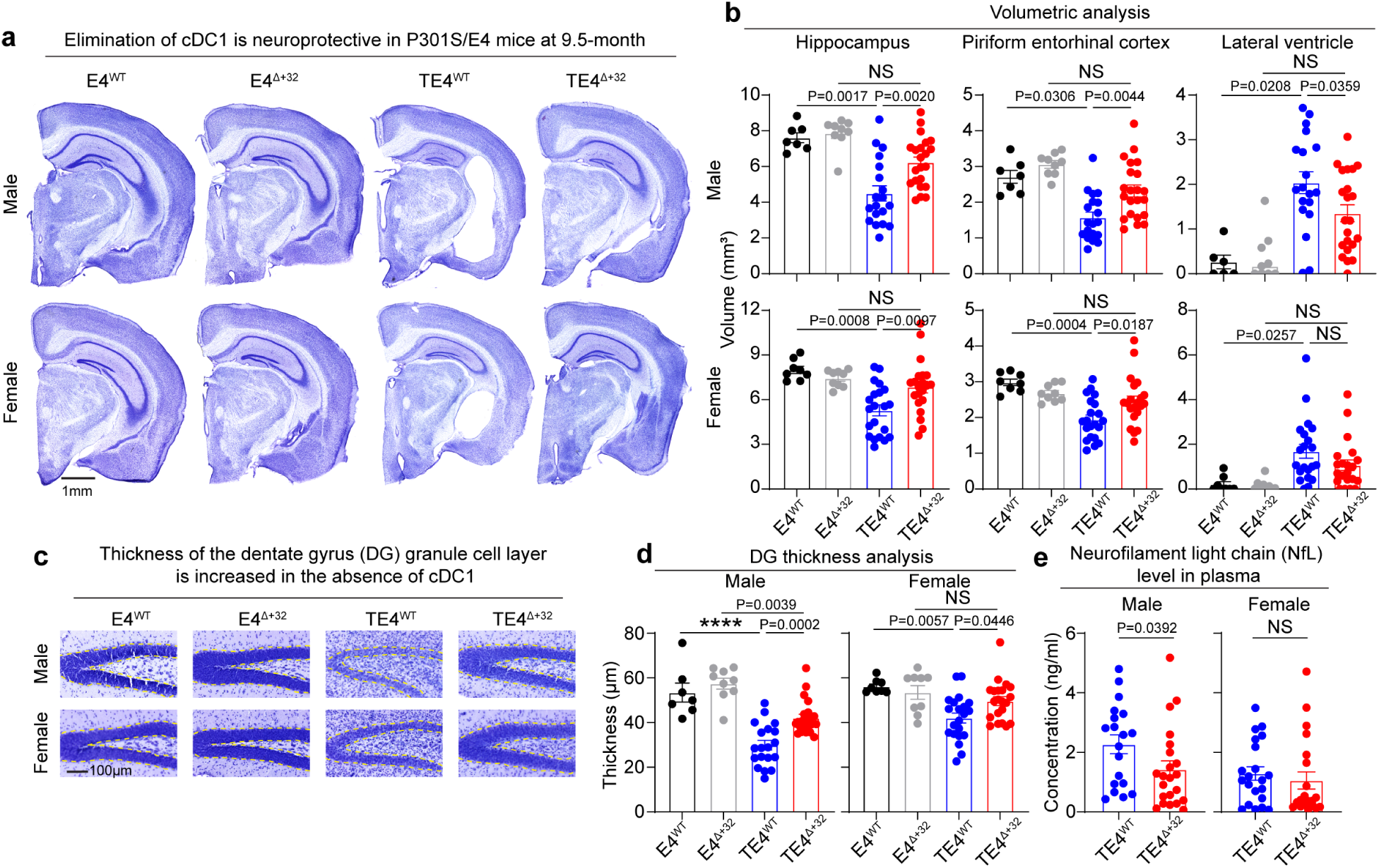
cDC1 deficient TE4 mice are protected against tau-mediated neurodegeneration. (**a**) Representative images of 9.5-month-old male (upper) and female (lower) E4^WT^, E4^Δ+32^, TE4^WT^, and TE4^Δ+32^ mice. (**b**) Volumetric analysis of the hippocampus, piriform entorhinal cortex and lateral ventricles of male (upper) and female (lower) E4^WT^ (n=7 for male and n=8 for female), E4^Δ+32^ (n=9 for both sexes), TE4^WT^ (n=19 for male and n=22 for female), and TE4 ^Δ+32^ (n=21 for both sexes) mice. (**c**) Representative images showing the thickness of the dentate gyrus (DG) granule cell layer in four strains of mice. (**d**) DG thickness analysis in four strains of mice. (**e**) Level of plasma neurofilament light chain (NfL) in male or female TE4^WT^ and TE4 ^Δ+32^ mice. Data are presented as mean values ± SEMs. Statistical significance was defined using two-way ANOVA with Šídák’s multiple comparisons test (**b** and **d**) and Mann-Whitney test, two-tailed (**e**). P value (actual value) was indicated in the figure. ****P<0.0001; NS, no significance.

### cDC1 deficiency does not strongly affect tau phosphorylation or aggregation

To test whether cDC1 deficiency affected tau pathology we performed immunohistochemical staining as well as quantitative ELISA to assess the level of soluble and insoluble tau and tau hyperphosphorylation (p-tau) at 9.5-months of age (Extended Data Fig. 4a). We did not observe a significant difference in phosphorylated tau (AT8) staining in the hippocampus between TE4^WT^ and TE4^Δ+32^ mice in either males or females (Extended Data Fig. 4b,c). The levels of total tau and p-tau were further analyzed in cortical tissues following sequential tissue extraction in RAB (soluble), RIPA (less soluble) and 70% formic acid (insoluble)^24^. Both male and female TE4^Δ+32^ mice presented higher level of total tau in RAB fraction (Extended Data Fig. 4d) but no significant difference in RIPA or formic acid fractions (Extended Data Fig. 4e,f). We found slight, but statistically lower levels of p-tau in both RAB and RIPA fractions but not in FA fractions in TE4^Δ+32^ compared to TE4^WT^ mice (Extended Data Fig. 4g-i). These data suggested that tau pathology was not markedly affected by cDC1 deficiency.

### cDC1 deficient TE4 mice exhibit reduced microglia and astrocyte reactivity

Microglial activation drives tau-mediated neurodegeneration and can be promoted by T cell infiltration into the brain^1^. We assessed hippocampal microglia immunostaining using IBA1, MHC-II (an IFNγ-responsive marker), and P2RY12 (a homeostatic microglial marker). Consistent with reduced neurodegeneration, immunostaining of IBA1 and MHC-II was significantly reduced in TE4^Δ+32^ compared to TE4^WT^ in male mice (Fig. 2a,b and Extended Data Fig. 5a). P2RY12 immunostaining was not significantly different in TE4^WT^ compared to TE4^Δ+32^ but immunostaining was increased in male E4^Δ+32^ compared to E4^WT^ (Fig. 2b and Extended Data Fig. 5a). Microglial staining for all three markers was not significantly different between female TE4^Δ+32^ and TE4^WT^ mice possibly due to the lower level of brain damage overall in female mice (Extended Data Fig. 5b-e). To assess astrocyte reactivity, we stained for glial fibrillary acidic protein (GFAP) and vimentin, two top elevated proteins enriched in reactive astrocytes in tauopathy mice^24,25^. cDC1 deficient male TE4^Δ+32^ mice exhibited significantly lower levels of both GFAP and vimentin hippocampal immunostaining than TE4^WT^ (Fig. 2c,d and Extended Data Fig. 5f). Furthermore, a robust co-colocalization of GFAP and vimentin can be detected in TE4^WT^ mice with greater brain atrophy but the overlap of the two proteins was scarce in TE4^Δ+32^ mice (Fig. 2e,f). GFAP immunostaining, but not vimentin, was also reduced in female TE4^Δ+32^ mice (Extended Data Fig. 5g-i). Overall, these results indicate that cDC1 deficiency reduces microglia and astrocyte reactivity.

**Fig. 2.**
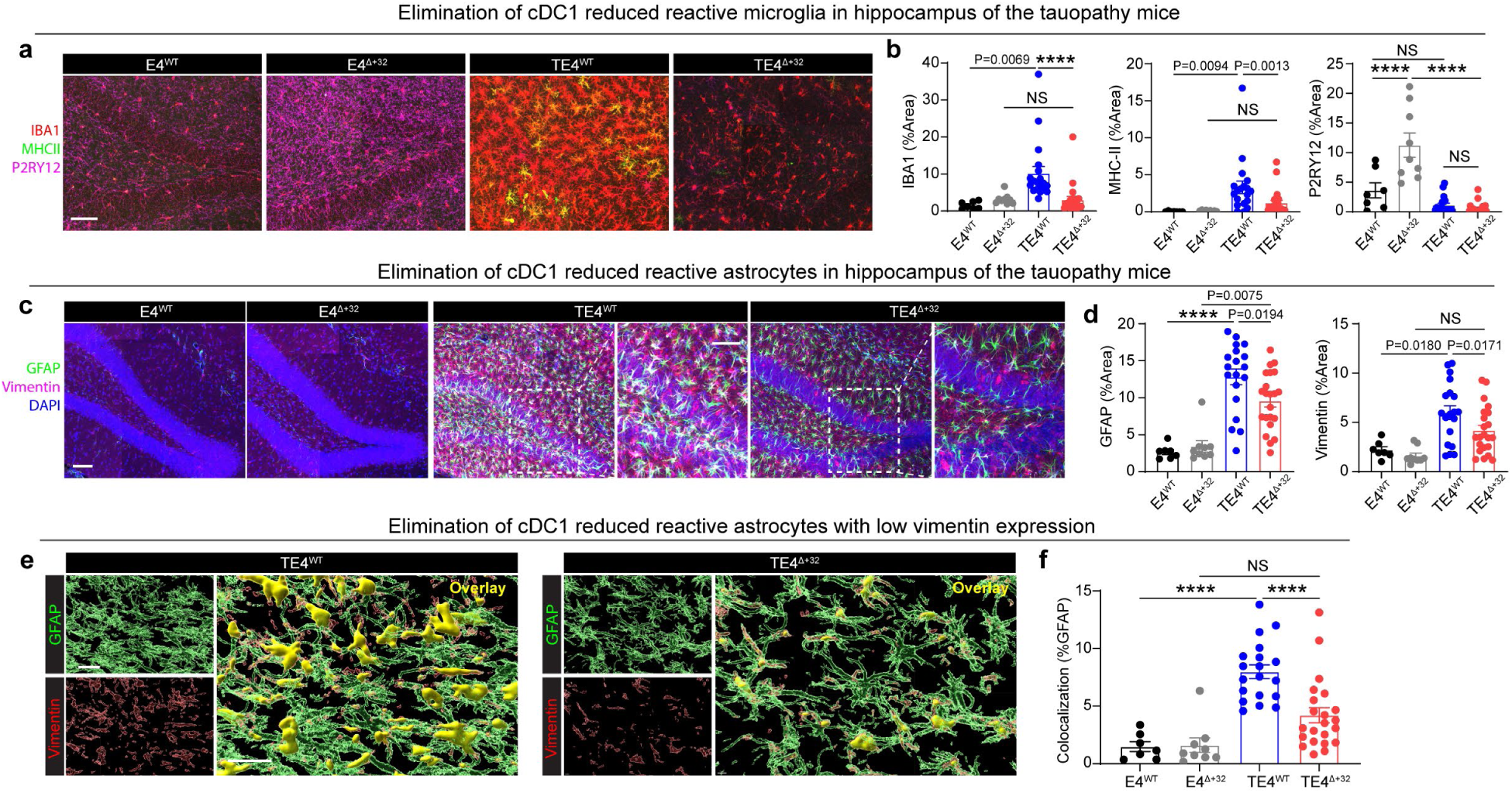
cDC1 deficient TE4 mice exhibit reduced reactive gliosis. (**a**) Representative immunofluorescent images of reactive microglia from four strains of the male mice at 9.5-month-old. The hippocampal sections were stained with IBA1 (Red), MHC-II (Green), and P2RY12 (Magenta). (**b**) Percent of the area analysis covered by IBA1 (Left), MHC-II (Middle), and P2RY12 (Right). (**c**) Representative immunofluorescent images of reactive astrocytes from four strains of the male mice at 9.5-month-old. The hippocampal sections were stained with Vimentin (Red), GFAP (Green), and DAPI (Blue). (**d**) Percent area covered by GFAP (Left) and Vimentin (Middle) staining. (**e**) Representative images showing the co-localization of Vimentin with GFAP from hippocampal sections from TE4^WT^ or TE4^Δ+32^ mice. (**f**) Percentage of Vimentin staining overlapping with GFAP staining from the sections from the indicated mice. n=7 for E4^WT^, n=9 for E4^Δ+32^, n=19 for TE4^WT^, and n=21 for TE4 ^Δ+32^ mice. Data are presented as mean values ± SEMs. Statistical significance was defined using two-way ANOVA with Šídák’s multiple comparisons test. P value (actual value) was indicated in the figure. Scale bar, 25μm. ****P<0.0001; NS, no significance.

### CD8^+^ T cells are selectively reduced in E4^Δ+32^ and TE4^Δ+32^ mice

To better understand the effect of cDC1 deficiency on lymphocyte populations, we performed single-cell RNA sequencing (scRNA-seq) on sorted live CD45^hi^ leukocytes from hippocampal and cortical tissues from 9.5-month-old E4^WT^, E4^Δ+32^, TE4^WT^, and TE4^Δ+32^ mice (Extended Data Fig. 6a,b). In E4^WT^ and E4^Δ+32^ mice, the frequency of CD45^hi^ leukocytes ranged between 0.4-1% of singlets, while, as described previously, leukocytes in TE4^WT^ increased approximately 4 to 5-fold to 2-4%^1^. However, cDC1-deficient TE4^Δ+32^ mice had reduced lymphocytes compared to TE4^WT^, albeit still elevated compared to E4^Δ+32^ (Extended Data Fig. 6c). Unbiased graph-based clustering of the brain infiltrating leukocytes revealed 11 cellular clusters with distinct gene signatures including conventional T cells, NK cells, NKT-like cells, B cells, plasmacytoid dendritic cells (pDC), innate-like lymphocytes (ILC2/3), and myeloid cells such as microglia (Fig. 3a,b and Extended Data Fig. 6d). The presence of leukocyte populations related to these clusters were validated by flow cytometry detecting various corresponding surface proteins or transcription factors. (Fig. 3b and Extended Data Fig. 7a-d).

**Fig. 3.**
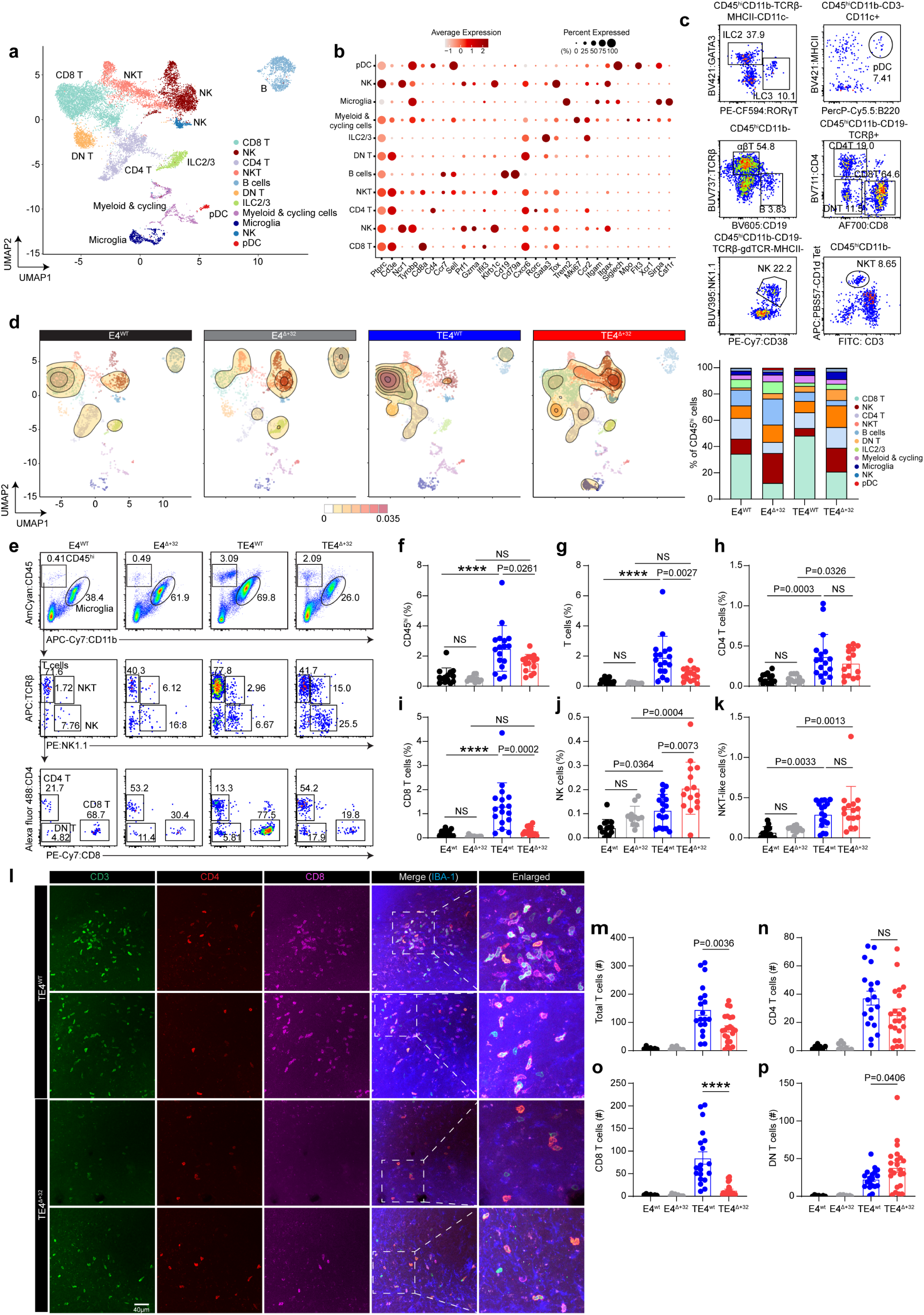
cDC1 deficiency specifically prevents CD8 T cells from accumulating in brain of tauopathy mice at 9.5 months of age. (**a**) UMAP plot shows individual clusters in CD45^hi^ leukocytes from brains of male E4^WT^, E4^Δ+32^, TE4^WT^, and TE4^Δ+32^ mice. (**b**) Dot plot identifies 11 clusters of leukocytes. (**c**) Representative flow cytometry plot confirms the identity of different leukocyte populations. (**d**) UMAP (left) plot reveals the population shift comparing the four strains of mice. Bar graph (right) shows the frequency of cells populating individual clusters. (**e**) Representative flow cytometry analysis identifying lymphocytes including conventional T cells (CD4 T cells and CD8 T cells), NK cells, and NKT cells in hippocampus and cortex of brains from 9.5-month-old male E4^WT^, E4^Δ+32^, TE4^WT^, and TE4^Δ+32^ mice. (**f-k**) Frequency of CD45^hi^ (**f**), αβ T cells (**g**), CD4 T cells (**h**), CD8 T cells (**i**), NK cells (**j**), and NKT cells (**k**) from hippocampus and cortex from 9.5-month-old male E4^WT^ (n=14), E4^Δ+32^(n=11), TE4^WT^(n=17), and TE4 ^Δ+32^(n=15) mice. (**l**) Representative immunofluorescence images of T cell accumulation in hippocampal regions of brains from the four strains of male mice. The hippocampal sections were stained with CD3 (Green), CD4 (Red), CD8 (Magenta), and IBA1 (Blue). Scale bar: 40μm. (**m-p**) Data summarizes the number of the CD3+ total T cells (**m**), CD8+ T cells (**n**), CD4+ T cells (**o**), and CD3+CD4-CD8- T cells (**p**) within the hippocampus. Male E4^WT^ (n=7), male E4^Δ+32^ (n=9), male TE4^WT^ (n=19), and male TE4 ^Δ+32^ (n=21). Data are presented as mean values ± SEMs. Statistical significance was defined using two-way ANOVA with Šídák’s multiple comparisons test (**f-k**) and unpaired student t test, two tailed (**m-p**). P value (actual value) was indicated in the figure. ****P<0.0001; NS, no significance.

When comparing cell-type compositions among groups we found that the proportion of CD8^+^ T cells was dramatically reduced in E4^Δ+32^ compared to E4^WT^ and in TE4^Δ+32^ compared to TE4^WT^ (Fig. 3d). There was a corresponding increase in the proportion of other lymphocyte populations in cDC1-deficent mice; however, CD4^-^CD8^-^ T cells, NK cells and NKT-like cells were particularly enriched in cDC1-deficient mice (Fig. 3d). A portion of CD4^-^CD8^-^ T cells consisted of invariant NKT cells which recognize lipid antigens^26^. The invariant NKT cells and NK cells are considered as the innate counterparts of adaptive immunity with antigen specificity or recapitulating features of cytotoxic CD8^+^ T cells^27–31^. Thus, one possibility is that these cells were compensating for the deficiency of brain infiltrating CD8^+^ T cells due to cDC1-deficiency.

To validate the findings from the scRNA-seq data, we performed flow cytometry analysis allowing us to quantitatively analyze the absolute changes in each population of interest. Around a five-fold increase in CD45^hi^ leukocyte infiltration can be detected in TE4^WT^ compared to E4^WT^ but TE4^Δ+32^ without cDC1 resulted in a roughly 40% reduction in the frequency the CD45^hi^ cells (Fig. 3e,f). Conventional T cells comprised approximately 70% of CD45^hi^ leukocytes in E4^WT^ and TE4^WT^ mice, but approximately 40% in E4^Δ+32^ and TE4^Δ+32^ mice resulting in an ∼50% reduction in T cells in cDC1 deficient mice (Fig. 3e,g). CD4^+^T cells and CD8^+^ T cells constituted the majority of the conventional T cells. CD4^+^ T cells were significantly increased with tau pathology but their frequency within the whole brain was not significantly different between TE4^WT^ and TE4^Δ+32^ mice (Fig. 3h), suggesting cDC1 did not affect the infiltration of CD4^+^ T cells into the brain during tauopathy. In stark contrast, the frequency of CD8^+^ T cells, which consisted of around 70% of conventional T cells in brain (Fig. 3e), was dramatically reduced in TE4^Δ+32^ compared to TE4^WT^ mice (Fig. 3i). These data suggest that the recruitment of CD8^+^ T cells in brain in the setting of tau-mediated neurodegeneration was largely abrogated by the absence of cDC1. Consistent with scRNAseq data, NK cells were increased in TE4^WT^ mice compared to E4^WT^, and their frequency was further increased in TE4^Δ+32^ mice (Fig. 3j). NKT-like cells [detected with TCRβ^int^ NK1.1^+^ (Fig. 3e)] were also increased in TE4^WT^, but we did not observe a further increase in TE4^Δ+32^ mice (Fig. 3k).

We next assessed the spatial localization and number of T cells within the hippocampus. We first verified that most of the CD3^+^ T cells were within the brain parenchyma and outside of the vasculature (Extended Data Fig. 8a). We then developed a tissue staining panel to simultaneously assess CD3, CD4, and CD8 expression. We also included IBA1 as marker for microglia. There was clear co-localization between either CD3 and CD4 or CD3 and CD8, corresponding to CD4^+^ T cells and CD8^+^ T cells respectively throughout the hippocampal region (Fig. 3l and Extended Data Fig. 8b). T cells were only rarely detected by immunostaining in E4^WT^ an E4^Δ+32^ brain (Extended Data Fig. 8c). In TE4^Δ+32^ mice, the number of total CD3^+^ T cells was only half of that in TE4^WT^ within the hippocampus (Fig. 3m). The density of the T cells within the hippocampal regions was negatively correlated with hippocampal damage (Extended Data Fig. 8d), suggesting again, that T cells contributed to neurodegeneration.

In TE4^Δ+32^ mice, the number of total CD3^+^ T cells was only half of that in TE4^WT^ within the hippocampus (averaged from two tissue sections of 50μm thickness) (Fig. 3m). The number of CD4^+^ T cells did not significantly differ between TE4^WT^ and TE4^Δ+32^ (Fig.3n). However, the number of CD8^+^ T cells was strongly reduced in TE4^Δ+32^ (Fig. 3o). CD4^-^CD8^-^ T cells were also significantly elevated in TE4^WT^ and further increased in TE4^Δ+32^ brains (Fig. 3p). These results revealed that the infiltration of CD8^+^ T cells was selectively abrogated by the absence of cDC1 and specifically implicated CD8^+^ T cells as promoting tau-mediated neurodegeneration.

### Reduced T cell activation in the brain of cDC1 deficient mice

We next further investigated the effect of cDC1-deficiency on specific T cell transcriptional states. Re-clustering the T cell population resulted in 14 transcriptionally different subsets (Fig. 4a). For CD4^+^ T cells, we identified memory T cells marked by intermediate expression of *IL7r*, *Cxcr6*, *Ccr7*, *Tcf7* etc. but broadly lacked the expression of *Sell* (Extended Data Fig. 9a,b). Within this group of memory CD4^+^ T cells, genes distinguished the central memory T cells (TCM) featured by the higher expression of *Ccr7*, *Klf2*, and effector memory T cells (TEM) featured by the higher expression of *Ctla2a*, *Nkg7*, *Ccl5* (Extended Data Fig. a,b). The memory CD4^+^ T cells also expressed a mixture of genes suggesting activation such as *Cd40lg*, *Tbx21*, *Icos*, etc. as well as inhibitory molecules such as *Tnfrsf18*, *Tnfrsf4*, *Ctla4*, *Izumo1r*, *Maf*, and *Hif1a*, imparting these cells an anergic signature with immunosuppressive activity (Extended Data Fig. 9a,b). We also identified regulatory T cells (Tregs) with distinctive *Foxp3* expression, and a subset of CD4^+^ T cells marked by high levels of *Il1rl1* (encoding ST2) and *Il2ra* (encoding CD25) (Extended Data Fig. 9a.b). For CD8^+^ T cells, the main subset was identified as an exhausted-like CD8^+^ T (Tex) cells with high expression of *Tox*, *Pdcd1*, and *Gzmk* (Fig. 4b), whose gene signatures recapitulated previously described age-associated T cells (Taa)^32–34^. A smaller Tox^int^Tcf7^int^ precursor exhausted-like T cell (Tpex) subset was also detected. Within the Tex cells, a population of cells distinctively expressed tissue resident genes such as *Igtae* (encoding CD103)^35^. A small subset of CD8^+^ T cells marked by activated NF-κB signaling with a high level of *Bcl3* expression likely representing memory precursor effector cells^36^ and differentiated memory CD8^+^ T cells with intermediate expression of *Tcf7*, *Cxcr6*, *Il7r*, etc.^37^ were found adjacent to the memory precursor cells in the UMAP plot, suggesting a close developmental relationship between the two subsets (Extended Data Fig. 9a,b). For both CD4^+^ and CD8^+^ T cells, naïve (*Sell*, *Ccr7*) and IFN-responsive signatures (*Isg15*, *Stat1*) were highly distinguishable to include both arms of T cells within the two subsets (Extended Data Fig. 9a,b). NKT-like cells exhibited distinctive expression of many NK cell receptors such as *Klrc1*, *Klra7*, *Klre1*, *Cd160* etc. as well as lack of either CD4 or CD8 expression (Fig. 4b and Extended Data Fig. 9a,b). Three different subsets of proliferating cells with high expression of genes involved in DNA replication^38^, anti-apoptosis^39^, and cytoskeleton regulation^40,41^ were also identified (Extended Data Fig. 9a,b).

**Fig. 4.**
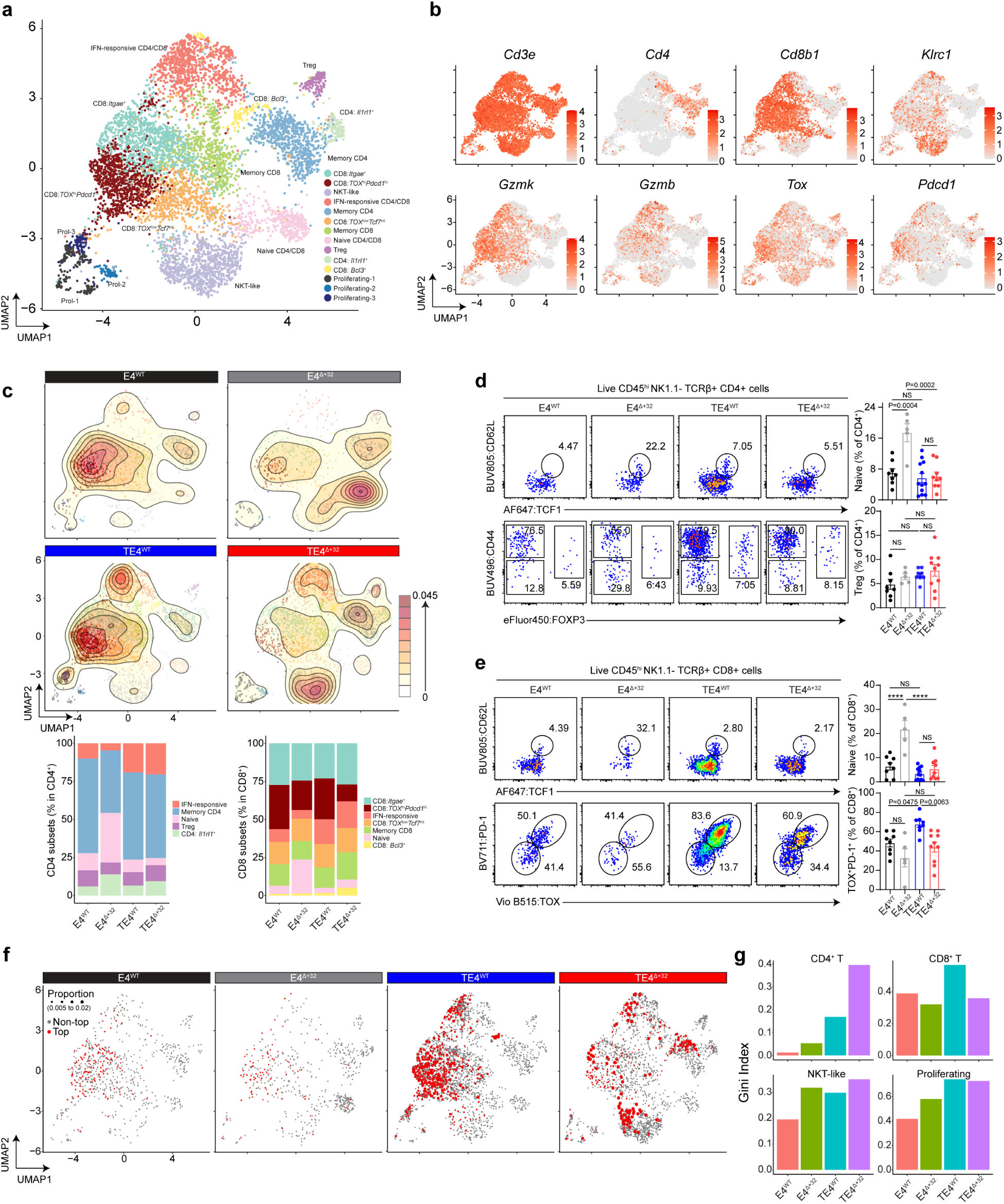
cDC1 deficiency leads to suboptimal activation of brain T cells and changes of TCR repertoire. (**a**) UMAP plot shows individual clusters in enriched CD3^+^ T cells from brains of E4^WT^, E4^Δ+32^, TE4^WT^, and TE4 ^Δ+32^ mice. (**b**) Feature plots show the expression of specific markers in clusters of cells. (**c**) Density plot (upper) shows the changes of the frequencies for each cluster and bar graph calculates the proportion of each cluster in CD4^+^ T cells (lower left) and CD8^+^ T cells (lower right). (**d**) Representative flow cytometry identifies naïve T cells (upper left) and Tregs (lower left) and frequencies of each population in CD4^+^ T cells among the four strains of male mice (right). (**e**) Representative flow cytometry identifies naïve T cells (upper left) and exhausted age associated T cells (lower left) and frequencies of each population in CD8^+^ T cells among the four strains of male mice (right). (**f**) UMAP plot shows the top T cell clonotypes in each strain of mice. Red dots represent top clones and size of the dots represented the proportion of the clonotypes among all T cells with paired TCR information (left). (**g**) Gini index bar plots of clonotypes in different cell types under different conditions. Higher Gini index indicates less even frequency of clonotypes and greater extent of clonal expansion. Data are presented as mean values ± SEMs. Statistical significance was defined using two-way ANOVA with Šídák’s multiple comparisons test (**d** and **e**). P value (actual value) was indicated in the figure. ****P<0.0001; NS, no significance.

Consistent with our previous observations^1^, the overall T cell number in TE4^WT^ was greater than in E4^WT^. Tex cells comprised the largest proportion of cells in TE4^WT^ and E4^WT^, but the proliferating cells and IFN-responsive cells (especially CD8^+^ T cells) were more abundant in TE4^WT^ mice (Fig. 4c and Extended Data Fig. 9c). This result suggested that tau pathology not only facilitated T cell infiltration into the brain but also led to local expansion and pro-inflammatory activity of CD8^+^ T cells. Activated CD8^+^ T cells, particularly Tex, were strongly reduced in E4^Δ+32^ mice while naïve cells were increased (Fig. 4c and Extended Data Fig. 9c). TE4^Δ+32^ mice also exhibited a strong reduction in activated CD8^+^ T cells, particularly Tex cell associated gene expression such as *Gzmk*, *Prf1*, *Tox*, and *Pdcd1* (Extended Data Fig. 9d). As a result, activated T cells within the brains of TE4^Δ+32^ mice were mostly comprised of activated CD4^+^ T cells including IFN-responsive cells, Tregs, as well as memory subsets and NKT-like cells (Fig. 4c and Extended Data Fig. 9c). Tregs are known to be immunosuppressive, and we observed that the many inhibitory molecules were highly expressed by the CD4^+^ memory T cell population (Extended Data Fig. 9a,b). These cells were enriched in TE4^Δ+32^ among the total T cell population. This may be largely due to the loss of CD8^+^ T cells because their proportion within the CD4^+^ T cells remained relatively unchanged (Fig. 4c). However, the enrichment of the immunosuppressive cells may still contribute to a level of protection.

We next validated scRNAseq observations using flow cytometry. We found that the proportions of naïve CD4^+^ and CD8^+^ T cells were significantly increased in E4^Δ+32^ mice and that the frequency of Tex cells was reduced by around 35% in TE4^Δ+32^ compared with that in TE4^WT^ mice (Fig. 4d,e and Extended Data Fig. 9e). Treg proportions did not significantly differ across groups (Fig. 4d). Overall, these findings suggest TE4^Δ+32^ mice exhibited decreased CD8^+^ T cell activation as reflected by the loss of IFN-responsive CD8^+^ T cells and Tex CD8^+^ T cells which may contribute to the protection against tau-mediated neurodegeneration.

### cDC1 deficiency decreases clonal expansion of CD8 T cells in TE4 mice

We next assessed the effect of cDC1 deficiency on clonal T cell expansion using paired TCR sequencing. Consistent with our previous findings, TE4^WT^ mice exhibited increased clonal expansion of CD4^+^ and CD8^+^ T cells compared to E4^WT^ mice (Fig. 4f). Tex and IFN-responsive CD8^+^ T cells from TE4^WT^ mice contained dominant T cell clones defined by the same paired TCR sequences (Fig. 4f and Extended Data Fig. 10a), suggesting that these cells recognized antigens presented in the brain. In contrast, clonally expanded T cells from TE4^Δ+32^ mice were enriched in memory/anergic CD4^+^ T cells and NKT-like cells with fewer represented clones in CD8^+^ T cells (Fig. 4f and Extended Data Fig. 10a). This was further validated by the observation that the most frequent clonotype in TE4^WT^ was found in CD8+ T cells while the most frequent clonotype in TE4^Δ+32^ mice expressed high levels of *Klrb1c*, suggesting an NKT-like cell origin (Extended Data Fig. 10b,c). Evaluation of the overall clonality of T cells using the Gini coefficient, we found decreased TCR diversity of CD4 T cells in TE4^WT^ and TE4^Δ+32^ mice. However, the Gini coefficient was lower for CD8^+^ T cells from TE4^Δ+32^ mice compared to TE4^WT^ mice (fig. 4g), suggesting a deficient CD8^+^ T cell response in the brain of mice without cDC1 cells. Interestingly, in both TE4^WT^ mice and TE4^Δ+32^ mice, the top frequent clonotypes were found in the proliferating cell subset, again suggesting a local expansion upon encountering cognate antigens (Extended Data Fig. 10d). Furthermore, most of the top proliferating clones (8 out of 20) were identified in TE4^WT^ mice and majority of them were CD8^+^ T cells whereas the top proliferating clones identified in TE4^Δ+32^ mice were found in NKT and CD4^+^ T cells (Extended Data Fig. 10d), suggesting in the absence of cDC1, the CD8^+^ T cells failed to optimally proliferate even after they infiltrated brain.

### cDC1-mediated cross-priming of CD8^+^ T cells with CNS antigens occurs in secondary lymphoid tissues outside CNS

cDC1 were rarely detected in brain parenchyma even with severe neurodegeneration (Extended Data Fig. 1f,g), yet they are critical for CD8^+^ T cell activation and neurodegeneration in the TE4 brain. We hypothesized that CNS antigen cross-presentation mediated by cDC1 is likely to occur outside the brain. To test this, we examined the ability of cDC1 to present a model antigen ovalbumin (Ova) to Ova specific CD8^+^ T cells (OT-1) *in vivo*. Ova was delivered exogenously in a cell-associated manner using cells derived from MHC-I triple KO (3xKO) (H-2K^b-/-^, H-2D^b-/-^, β2m^-/-^) mice (Extended Data Fig. 11a), which avoided autonomous Ova presentation by 3xKO cells. First, we intravenously delivered irradiated 3xKO splenocytes with osmotically loaded Ova (Extended Data Fig. 11b) to E4^WT^ or E4^Δ+32^ mice which were transferred with CellTrace Violet (CTV)-labeled naïve OT-1 cells one day prior. Consistent with previous reports, cell-associated Ova elicited OT-1 cell proliferation in the spleen from E4^WT^ mice but not E4^Δ+32^ mice, indicative of antigen cross-presentation deficits in the absence of cDC1 cells (Extended Data Fig. 11c).

Next, we aimed to mimic the cross-presentation of brain-derived antigens and define where antigens were presented. Ova-loaded 3xKO splenocytes or control splenocytes were injected into the hippocampus and cortex of E4^WT^ or E4^Δ+32^ mice through intracranial injection. OT-1 proliferation was examined (Extended Data Fig. 11d) in four different tissues: brain parenchyma, meninges, dCLN, and spleen. We did not detect any donor OT-1 cells present in the meninges or brain parenchyma under any conditions (Extended Data Fig. 11e) whereas OT-1 cells were found in dCLN and spleen. OT-1 cells underwent proliferation only in E4^WT^ provided with Ova-loaded splenocytes (Extended Data Fig. 11f). OT-1 proliferation was highest in dCLN (Extended Data Fig. 11g), suggesting that the presentation of brain-derived antigens occurred preferentially in dCLN and to some degree in the spleen of E4^WT^ mice.

We performed a similar experiment using cultured cortical neurons obtained from E15.5 3xKO pups virally transduced with AAV8 expressing a membrane-bound Ova (mOva) driven by a neuron-specific synapsin promoter (Fig. 5a). Again, transferred OT-1 cells were barely detectable in the meninges or brain in any condition (Fig. 5b) while OT-1 cells were present in secondary lymphoid tissues (dCLN and spleen) (Fig. 5c). Only in dCLN from E4^WT^ mice could we detect significant OT-1 proliferation (Fig. 5d). Together, these results suggest that cross-presentation of brain-derived antigens was mediated by cDC1, and this mainly occurred in the deep cervical lymph nodes, a major location of drainage of CSF and its constituents from the brain via the meningeal lymphatic system^42,43^.

**Fig. 5.**
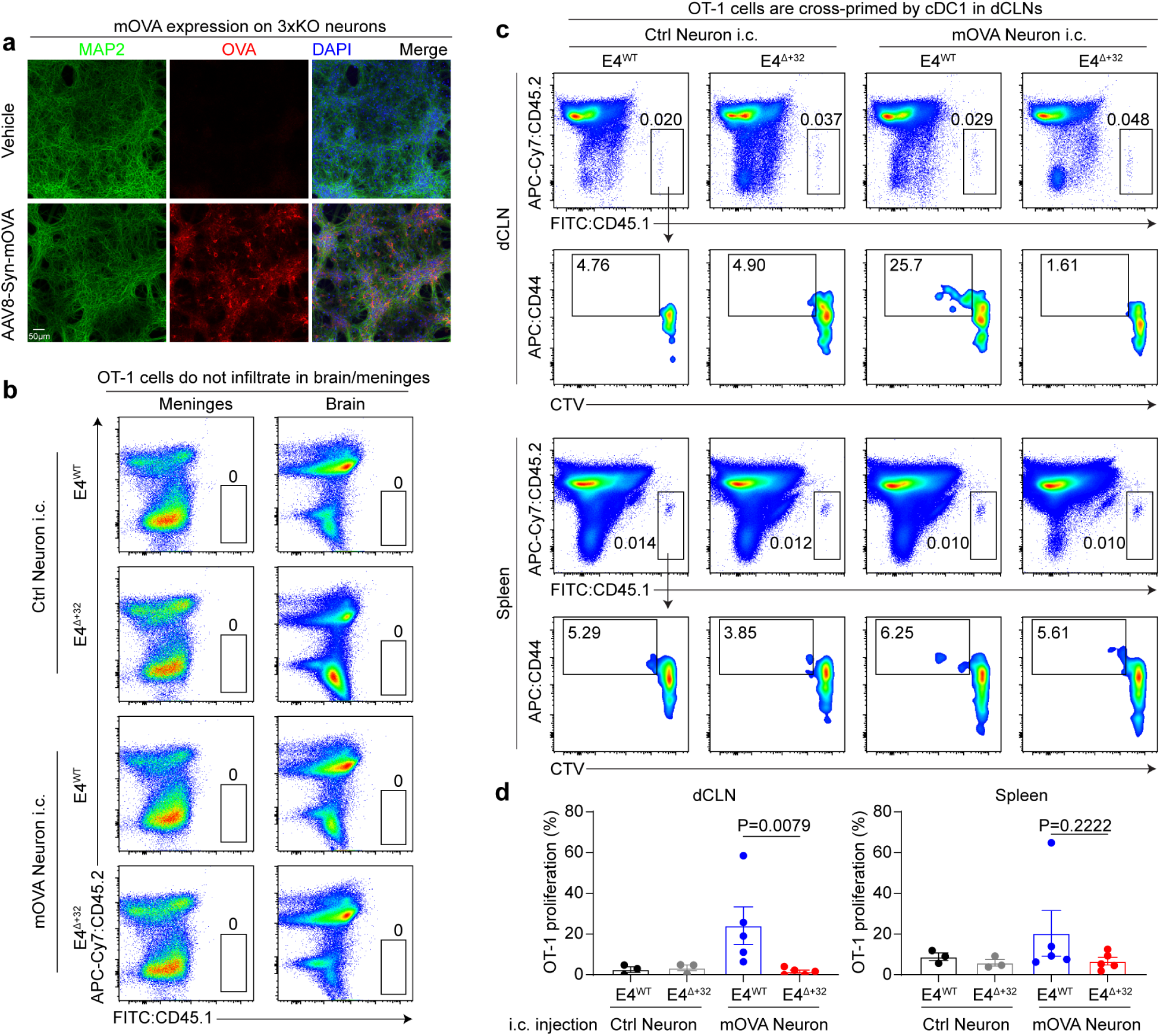
Cell-associated antigens derived from brain were cross-presented by cDC1 to CD8^+^ T cells in secondary lymphoid tissues outside of the CNS. (**a**) Representative immunofluorescent images of neurons derived from MHC-I 3xKO mice infected with or without AAV8-syn-mOVA virus. The cells were stained with MAP2 (Green), Ova (Red) and DAPI. (**b**) Representative flow cytometry shows transferred OT-I cells are not present in meninges or brain parenchyma. (**c**) Representative flow cytometry shows transferred OT-1 cells are present in dCLN and spleen and their proliferation rate under each condition. (**d**) Percentages of proliferated OT-I transferred into WT or Δ+32 mice injected with either control neurons or mOva expressing neurons into the brains. Data are presented as mean values ± SEMs. Statistical significance was defined using Mann-Whitney test, two-tailed. P value (actual value) was indicated in the figure.

Overall, we found that cDC1 cells were required for CD8^+^ T cell responses in the brain and that genetic ablation of cDC1 cells markedly protected against tau-mediated neurodegeneration. Our results suggest that targeting T cell priming by antigen presenting cells in the periphery should be further investigated as an approach to reduce tau-dependent neuroinflammation and degeneration.

## Methods

### Animals

P301S tau transgenic mice (PS19tg; Stock No. 008169, Jackson Laboratories) containing a transgene of the human P301S tau mutant driven by a mouse prion protein (Prnp) promoter were backcrossed to C57BL/6 mice (Stock No. 027, Charles River) for more than 10 generations. These mice were crossed with human APOE4 knock-in mice (C57BL/6 background) in which the endogenous mouse ApoE gene was replaced by a human APOE gene flanked by LoxP sites^44^ to generate a APOE4 homozygous P301S tau transgenic mice on a C57BL/6 background (abbreviated as TE4 mice). The TE4 mice were then crossed with *Irf8*+32^-/-^ mice with B57BL/6 background (Stock No. 032744, Jackson Laboratories) for two generations to generate E4^WT^, E4^Δ+32^, TE4^WT^, TE4^Δ+32^ mice. OT-1 mice (Stock No. 003831) containing transgenic inserts for mouse *Tcra-V2* and *Tcrb-V5* genes, encoding TCR receptors that recognize ovalbumin residues 257-264, were purchased from Jackson Laboratories. B6/CD45.1 mice (Stock No. 002014) carrying a differential pan leukocyte marker *Prprca* (known as CD45.1) were purchased from Jackson Laboratories. OT-1 mice were crossed with B6/CD45.1 mice for two generations to generate OT-1/CD45.1 mice for further experiments. MHC-I 3xKO (H-2K^b-/-^, H-2D^b-/-^, β2m^-/-^) mice were previously reported^45^ All mice were bred, maintained, and used in experiments in our specific pathogen-free (SPF) animal facility in accordance with the guidelines of the Division of Comparative Medicine under the protocols approved by the Institutional Animal Care Care and Use Committee (IACUC) at Washington University School of Medicine.

### Nest-building behavior

Two days before the sample collection from the mice, group-housed mice were housed individually. Pre-weighted nestlet (4 nestlets per cage) were provided in each cage. After an overnight housing (typically 16-18 hours), nest building behaviors were assessed and scored by an observer blinded to mouse genotype. A qualitative 5-point scale system was used and the score given based on percentage of remaining nesting material and shredded conditions. Briefly, score = 1, nest shredding <25%; 2, nest shredding 25 to 50%; 3, nest shredding 50 to 90%; 4, nest shredding >90%, but nest was not compacted yet; 5, complete nest built.

### Tissue collection and processing

For histopathology and biochemistry experiments, mice were euthanized at 285 days (around 9.5-month-old) of age by intraperitoneal injection of pentobarbital (200 mg/kg). Blood samples were collected in EDTA-treated tubes before cardiac perfusion with 3 U/ml heparin in cold Dulbecco’s PBS (DPBS, Gibco, Cat# 14040133). Blood samples were spun down (10 min, 2,000g, 4°C), and plasma was collected. Brains were carefully extracted, weighed, and cut into two hemispheres. The left hemisphere was collected for immunostaining/immunohistochemical analysis and immerse-fixed in 4% PFA overnight before being transferred to 30% sucrose and stored at 4°C until frozen in powdered dry ice and sectioned. Brains were cut coronally into 50μm sections on a freezing sliding microtome (Leica SM1020R) and stored in cryoprotectant solution (0.2 M DPBS, 15% sucrose, 33% ethylene glycol, 0.2μm filter for sterilization, made in house) at −20°C until used for experiments. The right hemisphere was dissected to isolate the hippocampus and the cortex for biochemical analysis, and the tissue was kept at −80°C until being analyzed.

For the flow cytometry experiments, mice were perfused using cold, heparinized DPBS. Spleens were collected and kept in cold DPBS for single cell preparation. Deep cervical lymph nodes (dCLN) were collected under a dissection scope in cold DPBS for single cell preparation. To isolate meninges and brains, the bottom part of the skull was carefully removed, exposing the undamaged brain parenchyma. Brains were carefully collected and kept in cold DPBS for single cell preparation and skullcaps with the adhered meninges were stored in cold RPMI1640 (Gibco, Cat# 11875085) supplemented with 2% FBS medium for single cell preparation. To isolate single cells from different tissues, the following procedures were conducted (all procedures were performed on ice unless noted otherwise) regarding each tissue type. For spleen, single cells were prepared by mechanically smashing the tissue through a 70μm pre-wet strainer (Corning) and red blood cells were removed by adding 1-2ml of red blood cell lysis buffer (8.3g NH4Cl, 1.0g KHCO3, 200μl of 0.5M EDTA diluted in 1L distilled H2O, filtered through 0.2μm filter for sterilization, made in house), agitating vigorously by pipetting and left at room temperature (RT) for 5min before neutralizing with 5-10 ml of DMEM media (Gibco, Cat# 11965084) supplemented with 10% FBS. For dCLN, the lymph nodes were digested with 250μg/ml of Collagenase B (Roche, Cat# 11088807001) and 30U/ml of DNase I (Roche, Cat# 11284932001) diluted in DPBS (∼1ml of digestion buffer for each sample) at 37°C for 30 min, agitating every 15min, and the enzymatic activity was neutralized by adding 5ml of cold MACS buffer (3% BSA, 2mM EDTA in DPBS, made in house) For dural meninges digestion, the meninges were digested in 2ml of digestion buffer in a 2ml eppendorf tube containing 0.5mg/ml collagenase P (Roche, Cat# 11213857001), 0.1mg/ml DNAse I (Roche, Cat# 11284932001) diluted in RPMI1640 supplemented with 2% FBS and incubated with shaking at RT for 1h. After digestion, the meninges were mechanically dissociated with pipetting up and down 15 times and the cell mixture was filtered through a pre-wet 100μm strainer (Corning) with cold RPMI1640 supplemented with 2% FBS and the filters were then washed at least twice to maximally recover the cells. Brains were dissected carefully to keep the hippocampal and cortical tissues. The tissues were put in 2ml of the digestion buffer containing 1mg/ml of the Collagenase IV (Sigma, Cat# C4-28-100MG) and 10 µg/ml DNase I ((Roche, Cat# 11284932001) and incubated in a 37°C incubator for 30 min (pipetting every 15 min). The dissociated cells were filtered through pre-wet 70μm strainer. Cells were spun down in a centrifuge at 500g X 5min except when noted otherwise (Cells isolated from meninges were spun down at 520g X 6min) and cell pellets were washed once with DPBS and supernatants were discarded. Cells containing myelin debris were resuspended with 40% isotonic Percoll (Sigma, Cat# P1644-1L). The Percoll stock solution was first diluted to isotonic “100%” Percoll by adding 9 parts of Percoll with 1 part of 10X HBSS solution (Gibco, Cat# 14065056) and the “100%” Percoll solution was further diluted to 40% by adding 2 parts of the “100%” Percoll solution and 3 parts of the 1X HBSS solution (Gibco, Cat# 14025076)] and spun at 1000g for 10 min at RT with deceleration of “2”. After centrifugation, the tubes were carefully taken out and the myelin layer (at top) as well as the supernatant was carefully removed, and the cell pellets were washed twice with DPBS and spun down for antibody labeling.

### Isolation and culture of BM progenitor cells

BM was harvested from the tibia and femurs. Single cells were prepared by passing through a 70μm pre-wet strainer (Corning) and red blood cells were removed by red blood cell lysis buffer (see “*<u>Tissue collection and processing</u>*“ section). To enrich CDP, pre-cDC1 and pre-cDC2 cells before subjected to cell sorting which was reported previously^46^, BM cells underwent negative selection by adding biotinylated antibody mastermix including antibodies specific to CD3, CD19, CD105, CD127, TER-119, Ly-6G, and B220, followed by depletion with MagniSort Streptavidin Negative Selection Beads (Invitrogen, Cat# MSNB-6002-74). The remaining lineage^−^ BM cells were then stained with fluorescent antibodies before sorting. CDPs were identified as lineage^−^SiglecH^-^CD117^int^CD135^+^CD115^+^MHCII^−^CD11c^−^ BM cells; pre-cDC1s were identified as lineage^−^SiglecH^-^CD117^int^CD135^+^MHCII^int–neg^CD11c^+^CD24^+^ BM cells; pre-cDC2s were identified as lineage^-^ SiglecH-CD117^int-low^CD135^+^MHCII^-^CD11C^+^CD115^+^ BM cells. For sequential gating/sorting strategies, see Extended Data Fig. 1.

Cells were sorted into Iscove’s Modified Dulbecco’s Medium (IMDM, Gibco, Cat# 12440053) supplemented with 10% FBS, 1% penicillin-streptomycin solution, 1% sodium pyruvate, 1% MEM non-essential amino acid, 1% l-glutamine solution, and 55 μM β-mercaptoethanol (complete IMDM). For dendritic cell cultures, sorted cells were cultured in complete IMDM supplemented with 100ng/ml recombinant Flt3L (PeproTech/Thermofisher, Cat# 250-31L-100UG) for 7 days. The cells were then subjected to flow cytometry analysis.

### Quantitative PCR (qPCR) analysis

The RNA from microglia cells sorted from the brain by FACS was isolated by using the PureLink RNA mini kit (Invitrogen, 12183020) and PureLink DNase (Invitrogen, 12185010) according to the manufacturer’s protocol. 50 nanograms of microglial RNA from each sample were used for quantification of gene expression using TaqMan Gene Expression Master Mix (Applied Biosystems, Cat# 4369016) per instruction. The following primer sets were used:

*Actb* Std DNA Primer 1: GAC TCA TCG TAC TCC TGC TTG

*Actb* Std DNA Primer 2: GAT TAC TGC TCT GGC TCC TAG

*Actb* PrimeTime Probe: /56-FAM/CTG GCC TCA /ZEN/CTG TCC ACC TTC C/3IABkFQ/

*Irf8 Std DNA Primer 1:* TGT CTC CCT CTT TAA ACT TCC C

*Irf8 Std DNA Primer 2:* GAA GAC CAT GTT CCG TAT CCC

*Irf8 PrimeTime Probe:* /56-FAM/ACC TCC TGA /ZEN/TTG TAA TCC TGC TTG CC/3IABkFQ/

### Brain volumetric analysis

The procedure started with cutting serial 50-um thick coronal sections collected from the rostral to caudal end of the forebrain of the left-hemisphere of each brain. Volumetric analysis of the hippocampus, EC/PC, and ventricle was performed via stereological methods by assessing sections spaced 300 μm starting from bregma −1.3 mm to bregma −3.1 mm (6–7 sections per mouse depending on the severity of brain atrophy). Sections were stained with 0.25% cresyl violet for 5 min at room temperature, dehydrated in different concentrations of ethanol (50%, 70%, 90%, 95% X 3, 100% X 2), cleared with xylene (2X) and then coverslipped with Cytoseal60 (Electron Microscopy Sciences, Cat# 18006). Slides were imaged with the Nanozoomer 2.0-HT system (Hamamatsu Photonics) at 20x magnification, and areas of interest were traced using the NDP Viewer software (Hamamatsu Photonics, U12388-01). The volume of region of interest was quantified using the following formula: volume = (sum of area) ∗ 0.3 mm. To quantify the thickness of the dentate gyrus granular cell layer, a scale was drawn perpendicular to the cell layer at two spots of the sections. Two sections, corresponding approximately to bregma coordinates −2.7 and −3.0 mm, were selected for quantification. The average thickness value for each mouse was determined.

### Plasma neurofilament light chain (NfL) concentration measurement

Plasma NfL concentration was measured with NF-Light Simoa Assay Advantage kit on a Samoa HD analyzer (Quanterix). The measurement was performed per the manufacturer’s instructions.

### Flow cytometry analysis and cell sorting

Single cells prepared from different tissues were incubated with MACS media supplemented with Fc-block (Purified anti-Mouse CD32/CD16 (Clone: 2.4G2), Leinco) and cells were stained with fluorescently labeled antibodies (all the antibodies including ones for intracellular staining used in this study were listed in supplemental data) at 4°C for 20 min. Cell were then washed and analyzed by flow cytometry or subjected to FACS sorting. For some experiments, intracellular transcription factors were stained. To do this, cells were firstly stained with surface antibodies at 4°C for 20 min. Then the cells were fixed and permeabilized using Foxp3/Transcription factor staining kit (eBioscience) per manufacturer’s instructions. Cells were stained with FOXP3, TCF1, TOX, T-bet, RORγT, GATA3 antibodies at RT for 30 min and then washed with permeabilization buffer before analyzed by flow cytometry. All conventional flow cytometry was done with BD FACSCanto II (BD Biosciences) or Cytek Aurora Spectrum Flow Cytometry (Cytek Biosciences, 5L UV-V-B-YG-R). To sort the brain infiltrating leukocytes for scRNA-seq analysis, single cells from brains were prepared as described above with one extra step during the 30 min digestion procedure. To avoid artifacts of cell activation induced during digestion, a previously described digestion buffer was used for cell dissociation^47^ (35260865). Briefly, the brain digestion buffer described above was supplemented with actinomycin D (5μg/ml, transcription inhibitor, Sigma), triptolide (10μM, transcription inhibitor, Sigma), and anisomycin (27.1μg/ml, translation inhibitor, Sigma). The single cell mixture was stained with antibodies specific for surface markers and propidium iodide at 4°C for 20 min and washed twice before subjected to cell sorting. To avoid cell death, the sorted cells were kept at 4°C in the sort chamber and collection rack controlled by the temperature control system throughout the sorting procedure and the collected cells were stored on ice after sort. To ensure enough cells for single cell sequencing, 3-4 mice from tau transgenic groups (TE4^WT^ and TE4^Δ+32^) and 5-8 mice from non-tau groups (E4^WT^ and E4^Δ+32^) were used. Time of the procedures was limited within 8 hours from sacrificing the mice to finishing the single cell library preparation. Around 1000 cells/μl (total of 10-30μl) of the cells were used for single cell library preparation. All the FACS sorting was done on BD FACSAria II sorter (BD Biosciences) or CytoFlex SRT (Beckman Coulter). For data analysis, flow cytometry data acquired from machines other than Cytek Aurora were exported as Fcs files and analyzed using FlowJo software (version 10.10.0, Tree Star). The data acquired from Cytek Aurora were unmixed using the SpectroFlo software first and then further analyzed using FlowJo software (version 10.10.0, Tree Star).

Microglia are defined as: CD45^int^,CD11b^+^

Brain ILCs are defined as: CD45^hi^,CD11b^-^,TCRβ^-^,CD11c^-^,MHC-II^-^,GATA3^+^(ILC2) or RORγT^+^(ILC3)

Brain pDC are defined as: CD45^hi^,CD11b^-^,CD11c^+^,B220^+^,MHC-II^+^

Brain αβT cells are defined as: CD45^hi^,CD11b^-^,TCRβ^+^,NK1.1^-^,CD4 (or CD8)^+^ Brain B cells are defined as: CD45^hi^,CD11b^-^,TCRβ^-^,CD19^+^

Brain γδT cells are defined as: CD45^hi^,CD11b^-^,TCRβ^-^,CD19^-^,MHC-II^-^,TCRγδ^+^

Brain NK cells are defined as: CD45^hi^,CD11b^-^,TCRβ^-^,NK1.1^+^

Brain NKT-like and iNKT cells are defined as: CD45^hi^,CD11b^-^,TCRβ^+^,NK1.1^+^ (NKT-like) or CD1d-PBS57 tetramer^+^ (iNKT cells)

cDC1 cells are defined as: CD45^+^,F4/80^-^.B220^-^,CD11c^+^,MHC-II^+^,XCR1^+^,Sirpα^-^cDC2 cells are defined as: CD45^+^,F4/80^-^.B220^-^,CD11c^+^,MHC-II^+^,XCR1^-^,Sirpα^+^

Brain naïve T cells are defined as: CD45^hi^,CD11b^-^,TCRβ^+^,NK1.1^-^,CD4 (or CD8)^+^,CD62L^+^,TCF1^+^

Treg cells are defined as: CD45^hi^,CD11b^-^,TCRβ^+^,NK1.1^-^,CD4^+^,CD44^int-pos^,FOXP3^+^

CD8 exhausted-like (or Taa) cells are defined as: CD45^hi^,CD11b^-^,TCRβ^+^,NK1.1^-^,CD8^+^,PD1^+^,TOX^+^

Taa cells can be further separated as: TCF1^+^SLAMF6^+^CD39^-^, TCF1^int^SLAMF6^-^CD39^+^, TCF1^-^SLAMF6^-^CD39^+^

### Brain protein extraction

RAB, RIPA, and 70% FA buffer were produced. The recipe for RAB buffer is 0.1 M MES, 0.5mM MgSO_4_, 1mM EGTA, 0.75 M NaCl, pH = 7.0. The recipe for RIPA buffer is 50mM Tris, 150mM NaCl, 0.5% sodium deoxycholate, 1% Triton X-100, 0.1% SDS, 5mM EDTA, and pH = 8.0. Complete Protease Inhibitor and phosSTOP were added freshly to RAB and RIPA buffer. Mouse cortical tissues were weighed and homogenized (around 10mg tissues were used for subsequent extraction) and sequentially homogenized using a homogenizer in cold RAB buffer at 10mg tissue/ 200μl buffer. Equal amount of RAB homogenates was transfer to centrifuge tubes and centrifuged 20min at 50,000 x g at 4°C. Supernatant was stored as RAB fraction at -80°C. Then equal amount of RIPA buffer was added to the pellet and sonicated for 1 min at 20% pulse, 1 s interval, followed by 20min of centrifugation at 50,000 x g at 4°C. Supernatant was collected as RIPA fraction and stored at -80°C. Then equal amount of 70% FA was added to the pellet. After the same procedure of sonication and centrifugation as above, supernatant was saved at FA fraction and stored at −80°C.

### Sandwich enzyme-linked immunosorbent assay (ELISA)

The levels of total human tau, and pTau in RAB, RIPA and 70% FA fractions were measured by sandwich ELISA and normalized to tissue weight. The coating antibodies for total human tau, and pTau were TAU-5 (mouse monoclonal, 20 μg/ml), and HJ14.5 (pThr181-tau, mouse monoclonal, 20 μg/ml, made in house), respectively. The capture antibodies total human tau, and pTau were HT7-biotinlyated (mouse monoclonal, 200 ng/ml) and AT8-biotinlyated (pSer202/pThr205-tau, mouse monoclonal, Thermo Fisher Scientific, MN1020B,, 300 ng/ml), respectively. Coating antibodies were applied at 50μl/well in 96-well half area clear flat bottom high-binding microplate (Corning, Car# 3690) overnight. The next day, plates were washed three times with TBS. Samples were diluted as following: for total tau: RAB, 1:5,000 dilution in ELISA buffer (0.5% BSA (RPI Research Products, Cat# 9048-46-8), 1X protease inhibitor in TBS); RIPA, 1:5,000 dilution in ELISA buffer; FA, first dilute with 1:20 in Tris buffer (pH 11), then dilute with 1:5 in ELISA buffer; for p-tau: RAB, 1:200 dilution in ELISA buffer; RIPA, 1:2,000 dilution in ELISA buffer; FA, first dilute with 1:20 in Tris buffer (pH 11), then dilute with 1:5 in ELISA buffer. Standards of recombinant tau (made in house) and p-tau peptides (synthesized) were used for quantification purpose. The diluted samples and standards were added in plates and incubated at 4°C overnight. The next day, the plates were washed three times with TBS, and developed with detection antibodies at RT for 2h with shaking. The plates were then washed three times with TBS and developed with Streptavidin-HRP at RT for 1h with shaking. Then the plates were washed, and the reaction was developed with adding 100μl of TMB substrate (Super Slow) (Sigma, Cat# T5569). The plates were read at 650nm by a plate reader every 5, 10, 15 min.

### Immunohistochemistry and imaging

The coronal sections from left hemispheres, with approximate coordinates −1.5 and −1.8 mm from bregma, were used for immunohistochemistry. Information on the antibodies can be found in Supplementary Table.

For AT8 staining, brain sections were washed in Tris-buffered saline (TBS) buffer three times followed by incubation in 0.3% hydrogen peroxide in TBS for 10 min at room temperature. After three washes in TBS, sections were blocked with 3% milk in TBS buffer with 0.25% Triton X-100 (TBSx) for 30 min at RT, followed by overnight incubation at 4°C with the biotinylated AT8 antibody (Invitrogen, MN1020B; 1:500) in blocking buffer. On the next day, brain sections were washed three times with TBS, followed by incubation in the VECTASTAIN Elite ABC-HRP solution (Vector Laboratories, PK-6100) at RT for 1 hour. Finally, after three washes with TBS, brain sections were developed and stained using the ImmPACT DAB EqV Substrate (Vector Laboratories, SK-1403). Tissues were mounted on the slides and dehydrated. Slides were then mounted with Cytoseal60 and scanned using a NanoZoomer microscope (Hamamatsu Photonics) at a 20X magnification. Images were extracted by using the NDP viewer software (Hamamatsu Photonics, U12388-01) and analyzed with ImageJ software (NIH, V1.54k).

### Immunofluorescence staining and imaging

Information on the antibodies can be found in Supplementary Table. For immunofluorescent staining of microglial and astrocytic markers, brain sections were washed in TBSx (TBS supplemented with 0.25% Triton X-100) for three times (10 mins/cycle). After washing, sections were blocked with a solution containing 10% normal donkey serum in TBSx for 1 hour at RT, followed by an overnight incubation at 4°C with primary antibodies [goat anti-IBA1 (Novus Biologicals, NB100-1028; 1:500), rat anti-MHCII (BioLegend, 107603; 1:1000), rabbit anti-P2RY12 (AnaSpec, AS-55043A; 1:1000), Alexa Flour 488-conjugated mouse anti-GFAP (Novus Biologicals, NBP2-33184AF488 (1:1000), and anti-Vimentin (Invitrogen, PA1-16759; 1:1000)] in a TBSx solution containing 2% normal donkey serum. On the next day, after three washes in TBSx (10 mins/cycle), brain sections were incubated with fluorescent-labeled secondary antibodies (Invitrogen, 1:1000) for 1 hour at RT. Sections were then incubated in DAPI solution (Invitrogen, 62248; 100 ng/mL) for nuclei staining, washed with TBSx, and mounted in ProLong Diamond Antifade mounting medium (Invitrogen, P36961). Images were obtained by using a Leica Stellaris 5 confocal microscope and further analyzed with ImageJ (NIH, v1.54k) and Imaris (Oxford Instruments, v10.2) software.

For immunofluorescent staining of T cells, brain sections were washed with TBSx for three times (10 mins/cycle). After washing, sections were blocked with a solution containing 10% normal donkey serum, 1% bovine serum albumin (BSA), and RecombiMAb anti-mouse CD16/CD32 FcR blocker (BioxCell, CP025; 1:400) in TBS buffer with 0.5% Triton X-100 for 2 hour at RT, followed by three days incubation at 4°C with primary antibodies [rat anti-CD3ε (Leinco Technologies, C2457 ; 1:400), human anti-CD4 (Miltenyi Biotec, 130-123-215; 1:50), rabbit anti-CD8α (Invitrogen, MA5-29682; 1: 200), and goat anti-IBA1 (Novus Biologicals, NB100-1028; 1:500)] in a solution containing 2% normal donkey serum and 0.2% BSA in TBS buffer with 0.1% Triton X-100. After three days incubation, brain sections were then washed in TBSx for three times (10 mins/cycle), followed by incubation with fluorescent-labeled secondary antibodies (Invitrogen, 1:500) for 2 hours at RT. Sections were washed with TBSx and mounted in ProLong Diamond Antifade mounting medium (Invitrogen, P36961). Images were obtained by using a Leica Stellaris 5 confocal microscope and further analyzed with ImageJ software (NIH, v1.54k).

### Immunocytochemistry

Information on the antibodies can be found in Supplementary Table. Primary cortical neurons cultured in the LAB-TEK chamber slides (Thermo Fisher Scientific, 177380) were fixed with 4% paraformaldehyde at RT for 10 mins. After three washes in TBSx buffer (10 mins/cycle), cortical neurons were blocked with a solution containing 5% normal donkey serum in TBSx for 1 hour in room temperature, followed by incubation with primary antibodies [chicken anti-MAP2 (Abcam, ab5392; 1:1000) and mouse anti-Ovalbumin (Invitrogen, MA5-15307; 1:1000)] at RT for another 1 hour. After three washes in TBSx (10 mins/cycle), cortical neurons were incubated with fluorescent-labeled secondary antibodies (Invitrogen, 1:1000) for 1 hour at room temperature. Cortical neurons were then incubated in DAPI solution (Invitrogen, 62248; 100 ng/mL) for nuclei staining, washed with TBSx, and mounted in ProLong Diamond Antifade mounting medium (Invitrogen, P36961). Finally, images were obtained by using a Leica Stellaris 5 confocal microscope.

### Image analysis

For AT8 coverage analysis, the hippocampus areas were selected from the two sections from each tissue. AT8 coverage analysis was performed by using ImageJ software (NIH, v1.54k) at a pre-set threshold parameter. For Microglia coverage analysis, fluorescent images from brain sections stained with microglial markers (IBA1, MHCII, and P2RY12) were acquired on a Leica Stellaris 5 confocal microscope at a 20X magnification and 1024*1024-pixel resolutions. For each mouse, z-stacked images from the region of hippocampus were taken from two separate brain sections. Microglia coverage analysis was performed by using ImageJ software (NIH, v1.54k). For astrocyte coverage and colocalization analysis, fluorescent images from brain sections stained with astrocytic markers (GFAP and Vimentin) were acquired on a Leica Stellaris 5 confocal microscope at a 20X magnification and 1024*1024-pixel resolution. For each mouse, z-stacked images from the region of hippocampus were taken from two separate brain sections. Astrocyte coverage analysis was performed by using ImageJ software (NIH, v1.54k). Level of colocalization on the astrocytic markers was analyzed using Imaris software (Oxford Instruments, v10.2). Astrocytes were traced and reconstructed using the Surface Tracer with a surface smoothing factor of 0.3 μm and an automated absolute intensity thresholding. Further surface-surface colocalization analysis were performed at a smoothing factor of 0.15. For T cell number quantification, fluorescent images from brain sections stained with T cell markers (CD3, CD4, and CD8) were acquired on a Leica Stellaris 5 confocal microscope at a 20X magnification and 1024*1024-pixel resolution. For each mouse, z-stacked images from the region of hippocampus were taken from two separate brain sections. T cell number quantification was performed by using ImageJ software (NIH, v1.54k). For this quantification, the cells were manually counted. To avoid bias, the images were blindly labeled and quantified by multiple researchers blinded to mouse type. Total T cells were counted by CD3^+^ cells; CD4^+^ T cells were counted as CD3^+^CD4^+^ T cells; CD8^+^ T cells were counted as CD3^+^CD8^+^ T cells; DN cells were counted as CD3^+^CD4^-^CD8^-^ cells; a very small portion of cells had all three markers and were counted as double positive (DP) cells.

### scRNA-seq and TCR-seq data analysis

scRNA-seq and TCR-seq data were aligned to mm10 using Cell Ranger V7.2.0. The filtered transcriptome alignment output of each sample was loaded into R as Seurat^48^ objects for quality control. Cells with mitochondrial RNA percentage > 5% were removed, and doublets were predicted and removed using DoubletFinder^49^ with parameters PCs = 1:30, pN = 0.25, pK = 0.09, assuming the doublet formation rate is 5%. Then samples were merged into one dataset and processed using log-normalization. Variable features were identified using the mean.var.plot method with a mean cutoff of 0.0125 ∼ 5, dispersion cutoff of > 0.5, and Tra/Trb/Igh/Igk genes removed from variable features. Reads are scaled by regressing out mitochondrial RNA percentage and number of features. All variable features were used for downstream analysis. PCA is performed using default parameters, and batch-effect is corrected using harmony^50^ by regressing over the samples. The top 40 harmony-corrected PCs were used for non-linear dimension reduction and clustering. Preliminary annotation was performed using resolution at 0.6, and one additional doublet cluster was removed based on overlapping gene expression signature. Lymphocytes were isolated for further processing and T cell subsets were recursively created until subsets with unanimous Cd8a or Cd4 expression were obtained. Top PCs for analysis of subsets were picked using ElbowPlot. Cell type (NKT, CD8T, CD4T, and proliferating) labels were added to each subset creation. Then, subsets were merged back into T cells and all-compartment objects. Artifact clusters with uniformly higher mitochondrial RNA percentages or non-biologically interpretable markers were removed during subset processing. Ribosomal RNA content was regressed during the scaling of variable features of the T cell subset. Markers were calculated using the FindAllMarkers function with only.pos=T.

TCR-seq data from four samples were loaded and processed using scRepertoire^51^ combineTCR function with removaNA=T and removeMulti=T. Clonotypes were called based on the amino acid sequences with clone frequency and proportion calculated for each sample. The relative abundance bar plot was made using the clonalHomeostasis function. Results from scRepertoire were merged into the T cell Seurat object using combineExpression with group.by = “sample” for further analysis and visualization. CloneSize was defined based on clonalProportion with the following levels: Small (1e-04 < X <= 0.001), Medium (0.001 < X <= 0.01), Large (0.01 < X <= 0.1), Hyperexpanded (0.1 < X <= 1). There were no hyperexpanded clones based on clonalProportion. Gini indices for each sample were calculated separately for each cell type using the Gini function from the DescTools package.

### Recombinant AAV production

The membrane bound OVA DNA sequence was PCR amplified from the pCI-neu-mOVA plasmid (Addgene, Plasmid# 25099) using the following primer pairs.

Forward primer mOVA-Fwd-SalI: ACG CGT CGA CGC CAC CAT GAT GGA TCA AGC CAG ATC AG

Reverse primer Ova-Rev-PmeI: AGC TTT GTT TAA ACT TAA GGG GAA ACA CAT CTG C

The amplified DNA was purified and digested by restriction endonuclease enzymes SalI (NEB, Cat# R3138S) and PmeI (NEB, Cat# R0560S) per instruction (https://nebcloner.neb.com/#!/redigest). The rAAV8 plasmid was provided by the Hope Center Viral Vectors Core at Washington University School of Medicine and digested with the same enzymes. The PCR products were inserted into the rAAV8 plasmid using T4

DNA ligase (NEB, Cat# M0202S) to make a rAAV8-syn-mOVA plasmid. The AAV viruses were prepared at the Hope Center Viral Vectors Core at Washington University School of Medicine. Briefly, the viruses were made within the packaging cell line, HEK293, which was maintained in DMEM medium, supplemented with 5% FBS, 100 units/ml penicillin, 100 μg/ml streptomycin in 37°C incubator with 5% CO2. The cells were plated at 30-40% confluence in CellSTACS (Corning) 24 h before transfection (70-80% confluence when transfection). 960μg total DNA (286μg of pAAV8, 448μg of pHelper, 226μg of AAV transfer plasmid) was transfected into HEK293 cells using polyethylenimine (PEI)-based method^52^. The cells were incubated at 37°C for 3 days before harvesting. The cells were lysed by three freeze/thaw cycles. The cell lysate was treated with 25 U/ml of Benzonaze at 37°C for 30 min and then purified by iodixanol gradient centrifugation. The eluate was washed 3 times with PBS containing 5% Sorbitol and concentrated with Vivaspin 20 100K concentrator (Sartorius Stedim, Bohemia, NY). Vector titer was determined by qPCR with primers and labeled probe targeting the ITR sequence^53^.

### Cortical neuron culture

Cortices were dissected from MHC-I 3xKO mice at approximately embryonic day 15.5 (E15.5) in Hibernate™-E Medium (Thermo Fisher Scientific). The dissected tissue was digested in 0.05% Trypsin-EDTA (Gibco) for 20 minutes at 37°C. Following digestion, the dissociated cortical neurons were washed twice with culture medium. The culture medium consisted of Neurobasal™ Medium (Thermo Fisher Scientific) supplemented with 1 μM 5-fluoro-2′-deoxyuridine, 1 μM uridine, B-27™ Supplement (Thermo Fisher Scientific), and 100 ng/mL 2.5S NGF (Harlan Bioproducts). Approximately 18 million cells were plated into 6-well tissue culture plates (Corning), which had been precoated with 0.1 mg/mL poly-D-lysine hydrobromide (Sigma-Aldrich) and 0.1 mg/mL mouse laminin (Thermo Fisher Scientific). The culture medium was replaced every 2–3 days, with half of the medium exchanged for fresh medium at each change.

### Intracranial injection

The irradiated cells were collected and diluted to 50,000 cells/μl for both splenocytes and primary neurons. Mice were anesthetized with isoflurane (induction: 5% isoflurane in oxygen; maintenance: 1-1.5% isoflurane in oxygen; flow rate: 0.3 L/min) and immobilized in a stereotactic frame (David Kopf Instruments). The resuspended cells (1 μL at each injection site) were injected into the dentate gyrus (coordinate: AP = -2.3 mm; ML = ±2 mm; and DV = -2 mm) and their overlying cortical region (coordinate: AP = -2.3 mm; ML = ±2 mm; and DV = -0.8 mm) bilaterally using a syringe (Hamilton, 7653-01) with removable 22s gauge (Hamilton, 7758-03) at a flow rate of 0.5 μL/min. Mice were allowed to recover on a 37°C heating pad and monitored for the first 72 hours after surgery.

### Antigen cross-presentation assays

8-12-week-old OT-1/CD45.1 mice were euthanized. Spleen and lymph nodes from multiple locations were harvested and the tissues gently smashed through a pre-wet 70μm strainer to dissociate cells (See “*<u>Flow cytometry analysis and cell sorting</u>*” section). Red blood cells were lysed by red blood cell lysis buffer and a single cell mixture was made. To enrich the T cells, cells underwent negative selection by adding biotinylated antibody mastermix including antibodies specific to CD19, B220, NK1.1, Ly6G, CD4, MHC-II, Ter-119. After incubating on ice for 20 min, cells were washed with MACS buffer and resuspended in MACS buffer at concentration of 10^7^ cells/100μl. Then 10μl/10^7^ cells of MagniSort Streptavidin Negative Selection Beads (Invitrogen, Cat# MSNB-6002-74) were added and incubated at RT for 5min. Then the cell/beads mixture was filled with MACS buffer to 5ml and place onto a magnet for 5min. The supernatant containing enriched cells was collected, washed, and stained with antibodies (listed in supplementary data) for cell sorting. Cells were sorted as streptavidin^-^B220^-^CD4^-^CD8^+^Vα2^+^Vβ5^+^CD62L^+^CD44^low^ cells, washed with DPBS, and stained with 1μM (final concentration) CellTrace Violet (Invitrogen, Cat# C34557) at 37°C for 15min. Staining was neutralized by adding 25ml of IMDM (Gibco, Cat# 12440053) supplemented with 10% FBS. Cells were counted and resuspended in DPBS at 2.5X10^6^ cells/ml. Recipient mice were intravenously transferred 5X10^5^ cells in 200μl. The next day, mice received antigens in three different conditions. Condition 1, for OVA-splenocytes, MHC-I 3xKO splenocytes were osmotically loaded with 10mg/ml soluble ovalbumin (Worthington Biochemical Corporation, Cat# LS003048), irradiated at 1,350rad, and 500,000 cells were injected intravenously into mice that had been injected with 500,000 CTV-labeled OT-1 T cells one day prior. After 3 days, spleens were harvested, mashed, and analyzed for CTV dilution of OT-1 cells. Condition 2, OVA-splenocytes, MHC-I 3xKO splenocytes were osmotically loaded with 10mg/ml soluble ovalbumin (Worthington Biochemical Corporation, Cat# LS003048), irradiated at 1,350rad, and 200,000 cells were injected intracranially at four different locations (See “*<u>Intracranial injection”</u>* section) into mice that had been injected with 500,000 CTV-labeled OT-1 T cells one day prior. After 5 days, brains, meninges, dCLNs, and spleens were harvested, mashed/digested, and analyzed for CTV dilution of OT-1 cells. Condition 3, neuron cultures were infected with AAV8-Syn-mOVA with MOI of ∼500 seven days after neurons were seeded. Three days after infection, cells were gently trypsinized and collected, irradiated at 3,000rad, and 200,000 cells were injected intracranially at four different locations (See “*<u>Intracranial injection”</u>* section) into mice that had been injected with 500,000 CTV-labeled OT-1 T cells one day prior. After 5 days, brains, meninges, dCLNs, and spleens were harvested, mashed/digested, and analyzed for CTV dilution of OT-1 cells.

### Statistics

All data were expressed as mean values ± SEMs. Statistical analysis was performed with GraphPad Prism 10. Means between two groups were compared with a two-tailed, unpaired Student’s t test or two-tailed, Mann-Whitney test. Two-way ANOVAs were used to analyze between-subjects designs with two variable factors. Tukey was used for post hoc pairwise comparisons. Fisher’s exact test was used to analyze probability distributions. The strength of the linear relationship between two different variables was analyzed using Pearson’s correlation. The null hypothesis was rejected at the P < 0.05 level. Statistical significance was taken as *P < 0.05, **P < 0.01, ***P < 0.001, and ****P < 0.0001. NS, P > 0.05, suggesting no statistical significance. The actual values were all provided in the figures except for ****P < 0.0001. NS, P > 0.05, which showed as “****” or “NS”.

## Data availability

Single-cell RNA sequencing (GEO: GSE279315) data were deposited in the National Center for Biotechnology Information Gene Expression Omnibus database with the access token: admzyaesvbwzpkr. All the data generated in this study are provided in the Supplementary Tables/Source Data file. Source data are provided in this paper.

## Code availability

Consolidated R codes for scRNA-seq bioinformatic analysis and visualization are deposited at 10.5281/zenodo.13913854.

## Acknowledgements

We thank the Genome Technology Access Center in the Department of Genetics at Washington University School of Medicine for technical support and assistance with data acquisition, processing, and initial analysis of the transcriptomic data. We thank the technical support by the Hope Center Alafi Neuroimaging, Viral Vectors, and Flow Cytometry Cores at Washington University School of Medicine. We thank all members of the Holtzman laboratory for the discussions and for providing technical support on many aspects of the study. This work was supported by a Carol and Gene Ludwig Award for Neurodegeneration Research (D.M.H.), National Institute of Health grant AG085374 (D.M.H.), NS090934 (D.M.H), the GHR Foundation (D.M.H.), the JPB Foundation (D.M.H.), Cure Alzheimer’s Fund (D.M.H.), Rainwater Charitable Foundation (D.M.H.), Alzheimer’s Association Research Fellowship AARF -23-1142708 (to H.H.), and NIH grant R01-AI129545 (to W.M.Y.).

## Contributions

H.H., J.D.U., and D.M.H. designed the study. H.H. and P.B-C.L. performed most of the experiments including tissue collection, cell sorting, flow cytometry analysis, most of the immunofluorescent staining, ELISA, antigen presentation and data analysis. C.Z. analyzed scRNA-seq data with advice from M.N.A.. H.H. and P.S. maintained mouse colony and related mouse husbandry. H.H. and Y.L. performed T cell immunofluorescent staining. H.J. performed AT8 immunohistochemical staining. P.S. and J.N. performed immunofluorescent staining to evaluate reactive astrocytes. R.A.O. and K.M.M provided *Irf8*+32^-/-^ mice and R.A.O. assisted the experiments related to antigen presentation. P.B-C.L. and W.T. performed *ex vivo* cortical neuron culture. S.L. and W.M.Y. provided MHC-I triple KO mice. S.L. assisted the flow cytometry analysis. H.H., J.D.U., and D.M.H. wrote the manuscript with contributions from all authors.

