## Supplementary figures and images for "CD8^+^ T cells are primed by cDC1 and exacerbate tau-mediated neurodegeneration"

### Supplemental Fig1

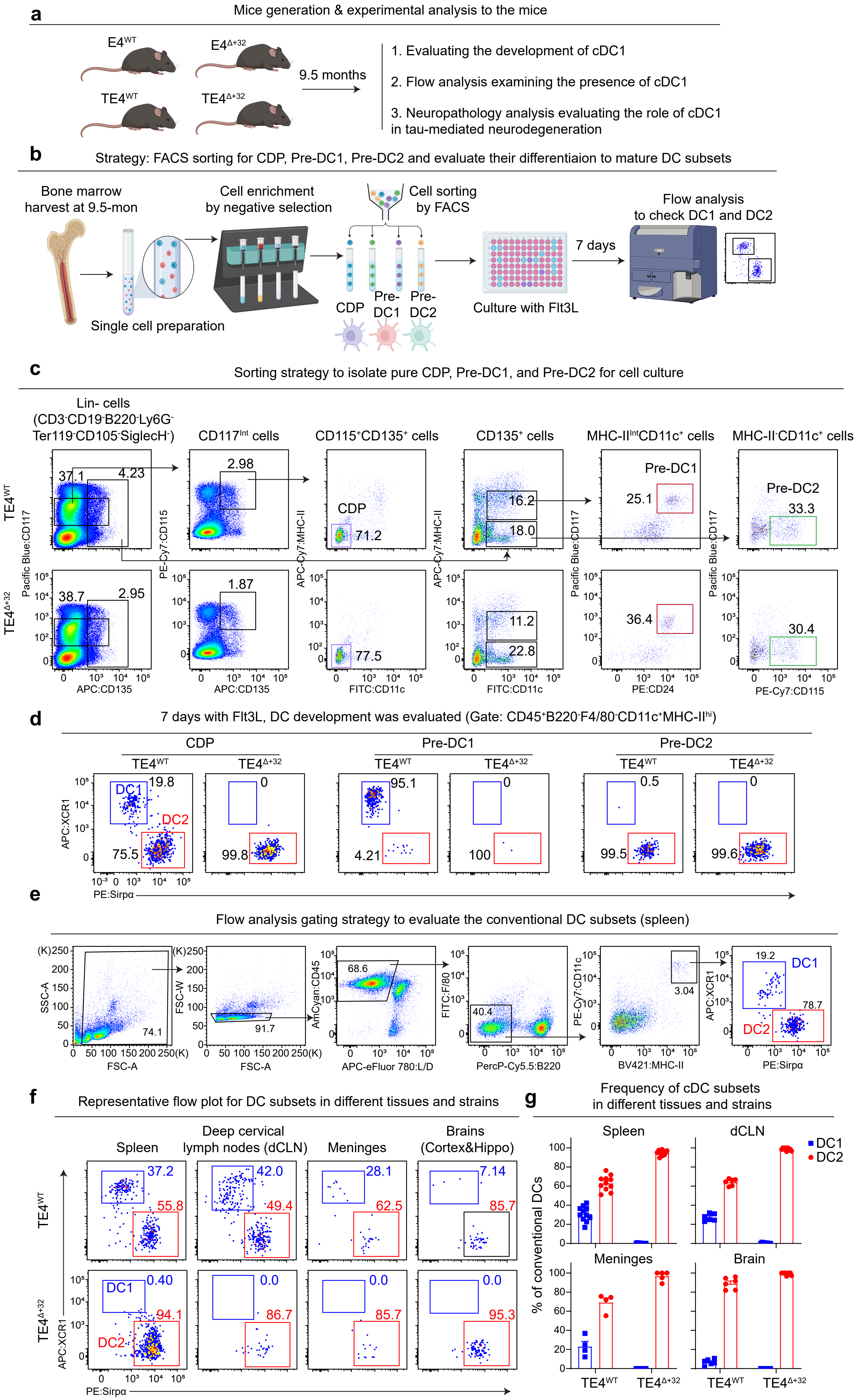

### Supplemental Fig2

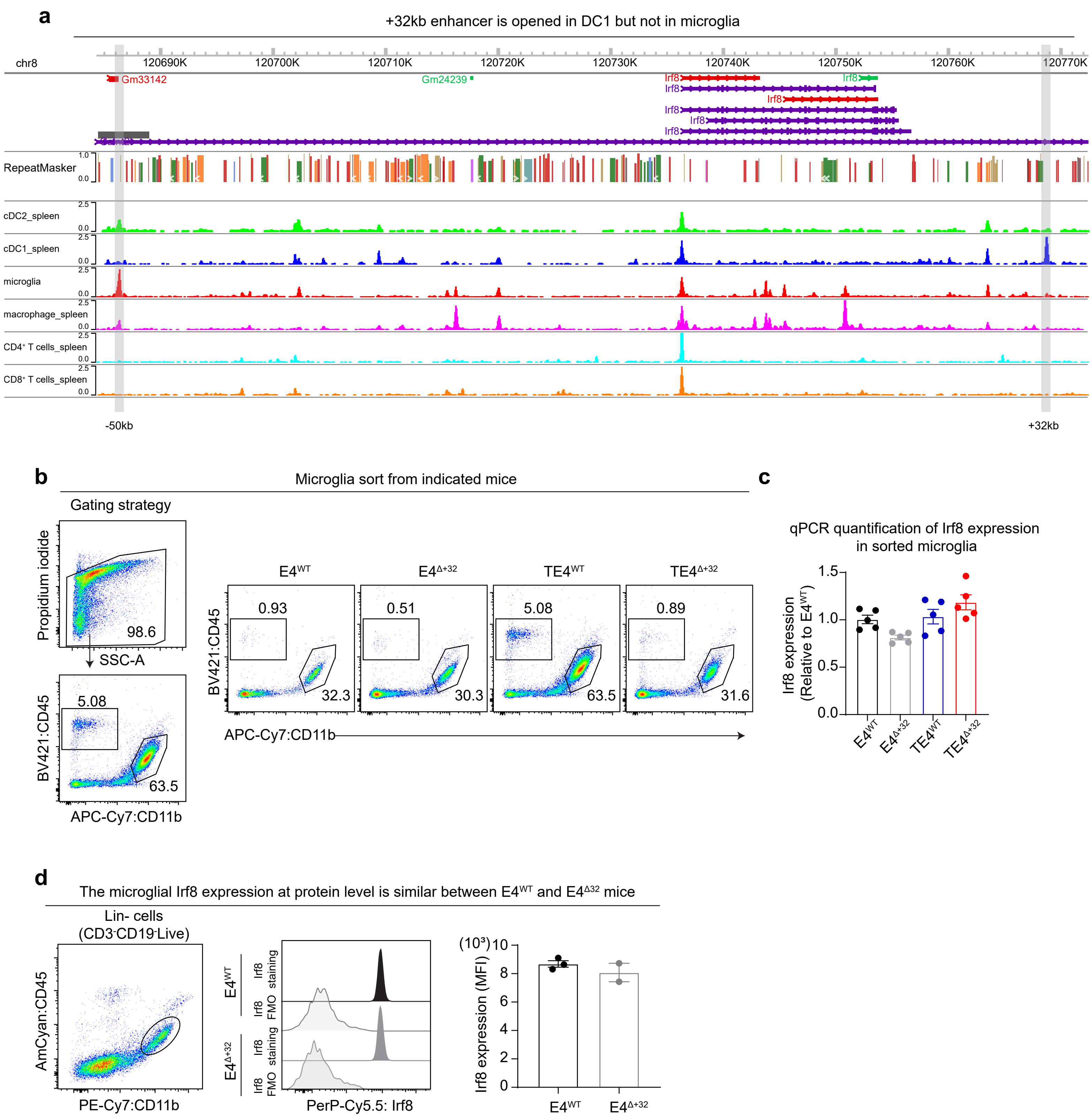

### Supplemental Fig3

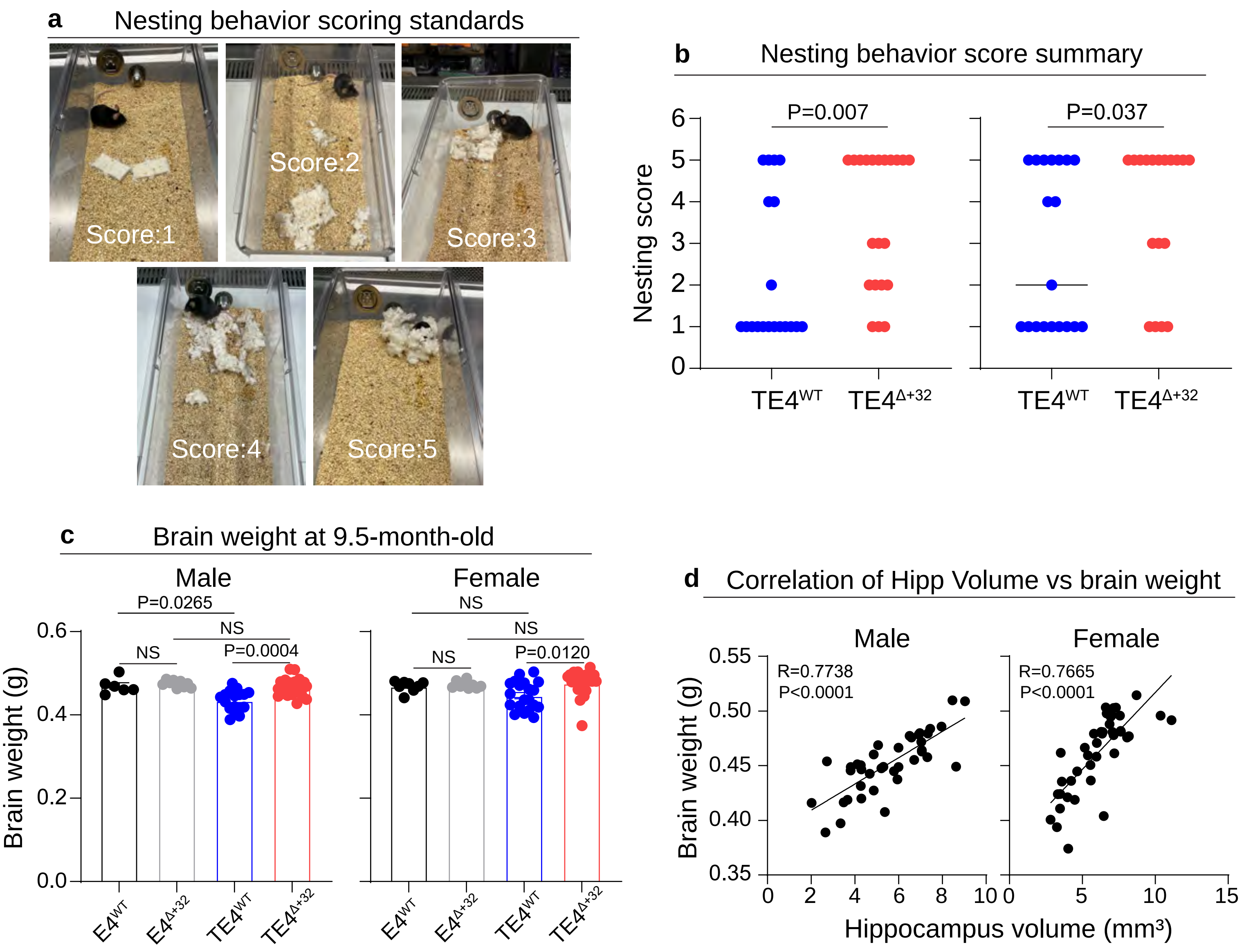

### Supplemental Fig4

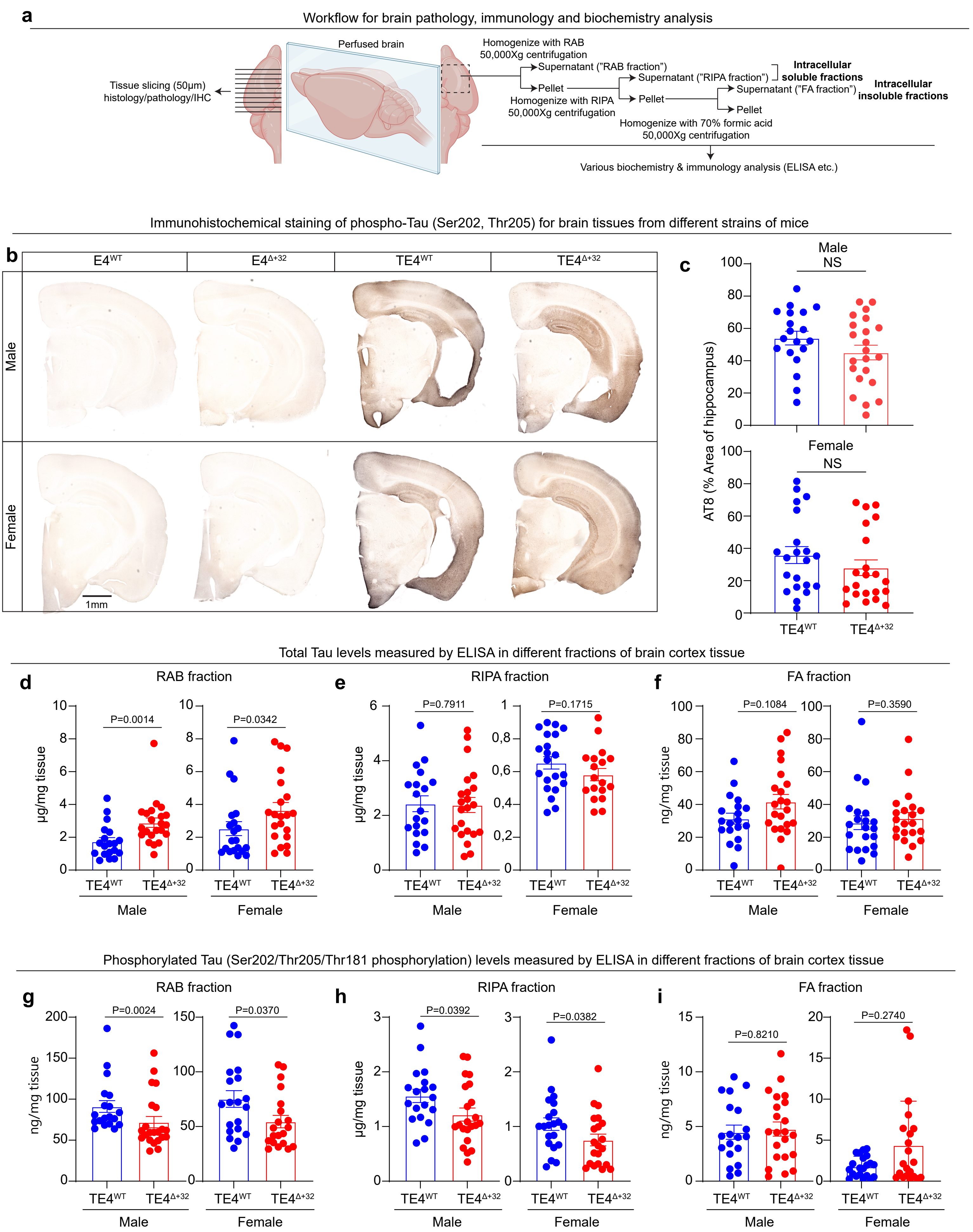

### Supplemental Fig5

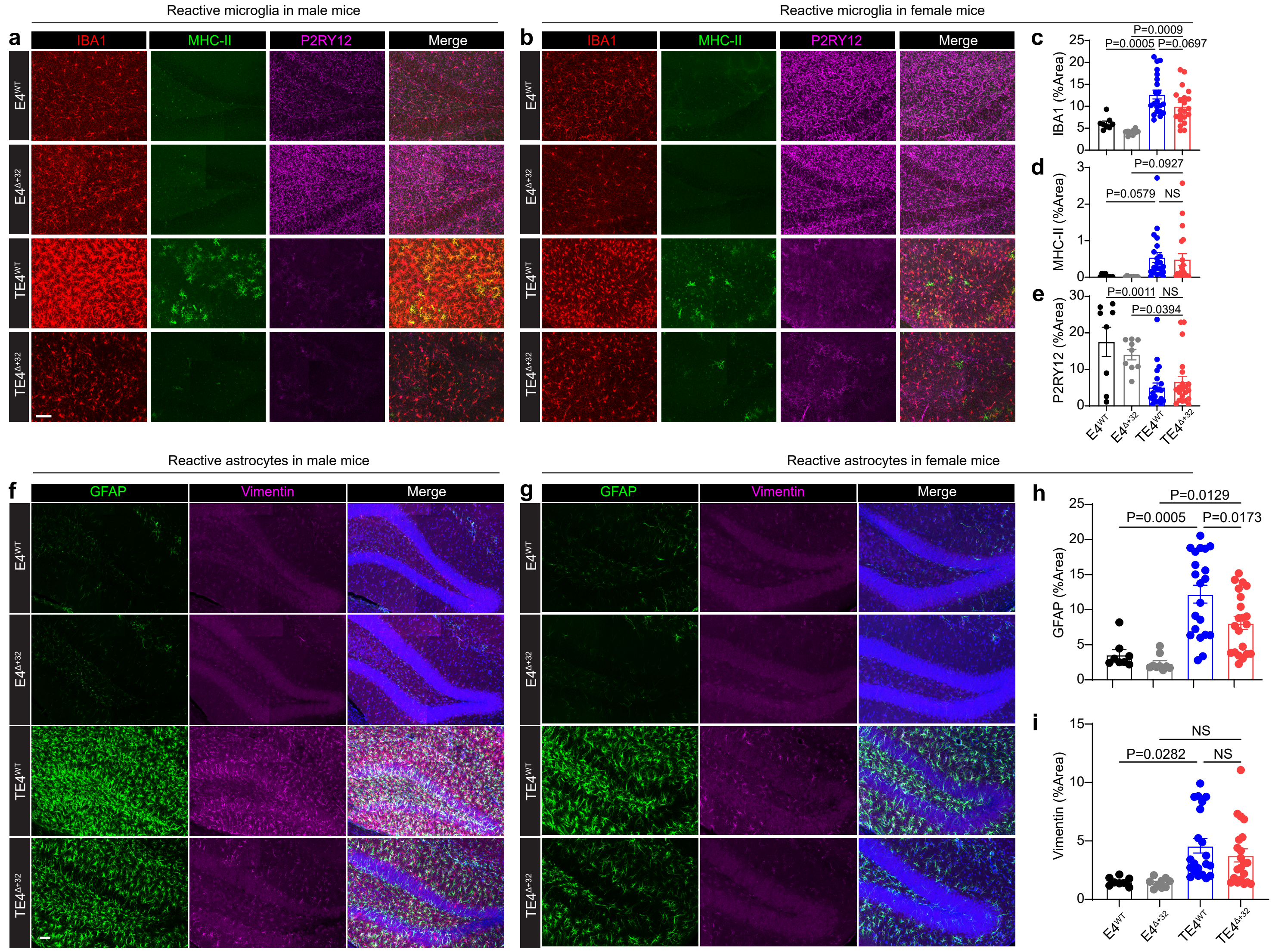

### Supplemental Fig6

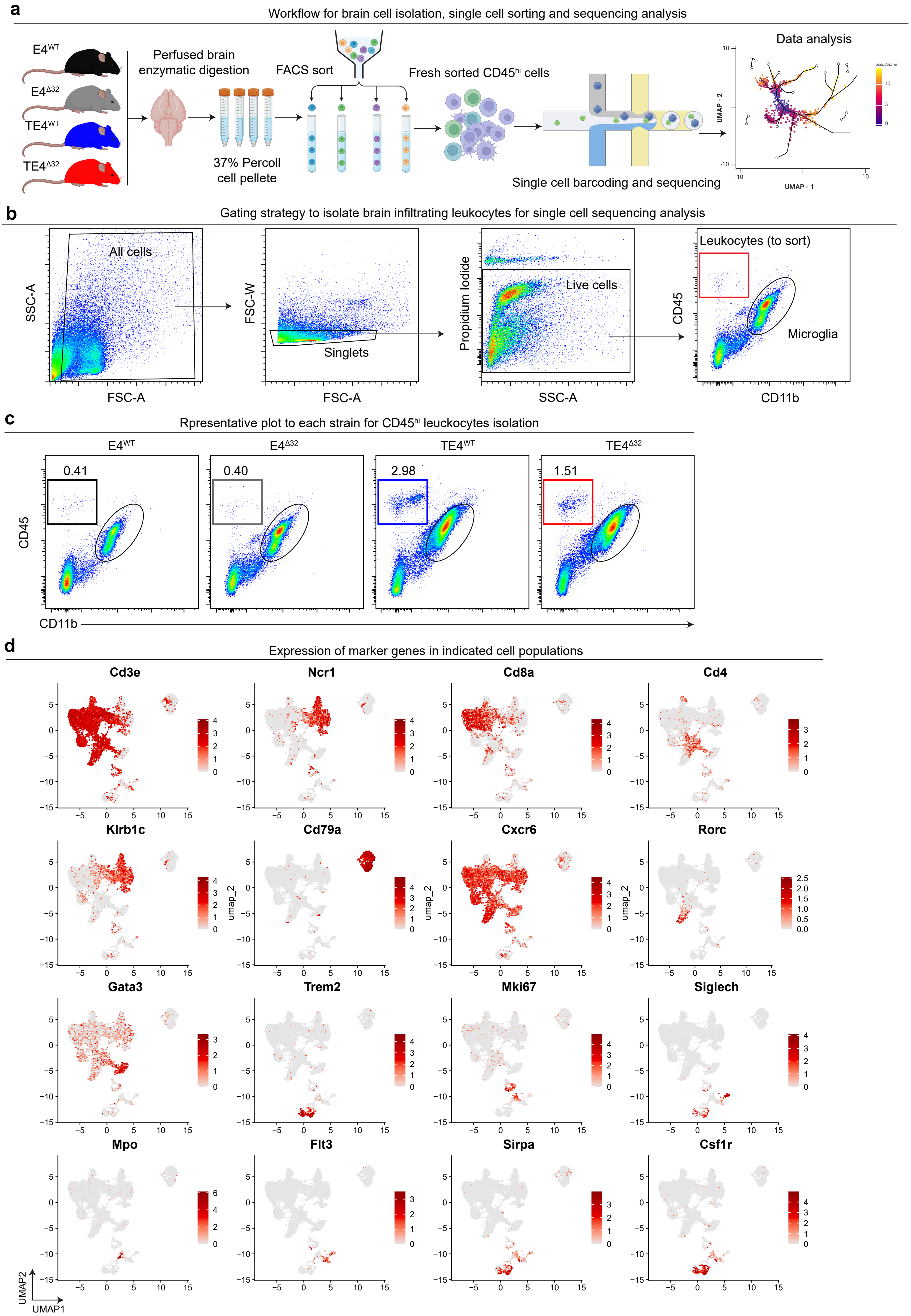

### Supplemental Fig7

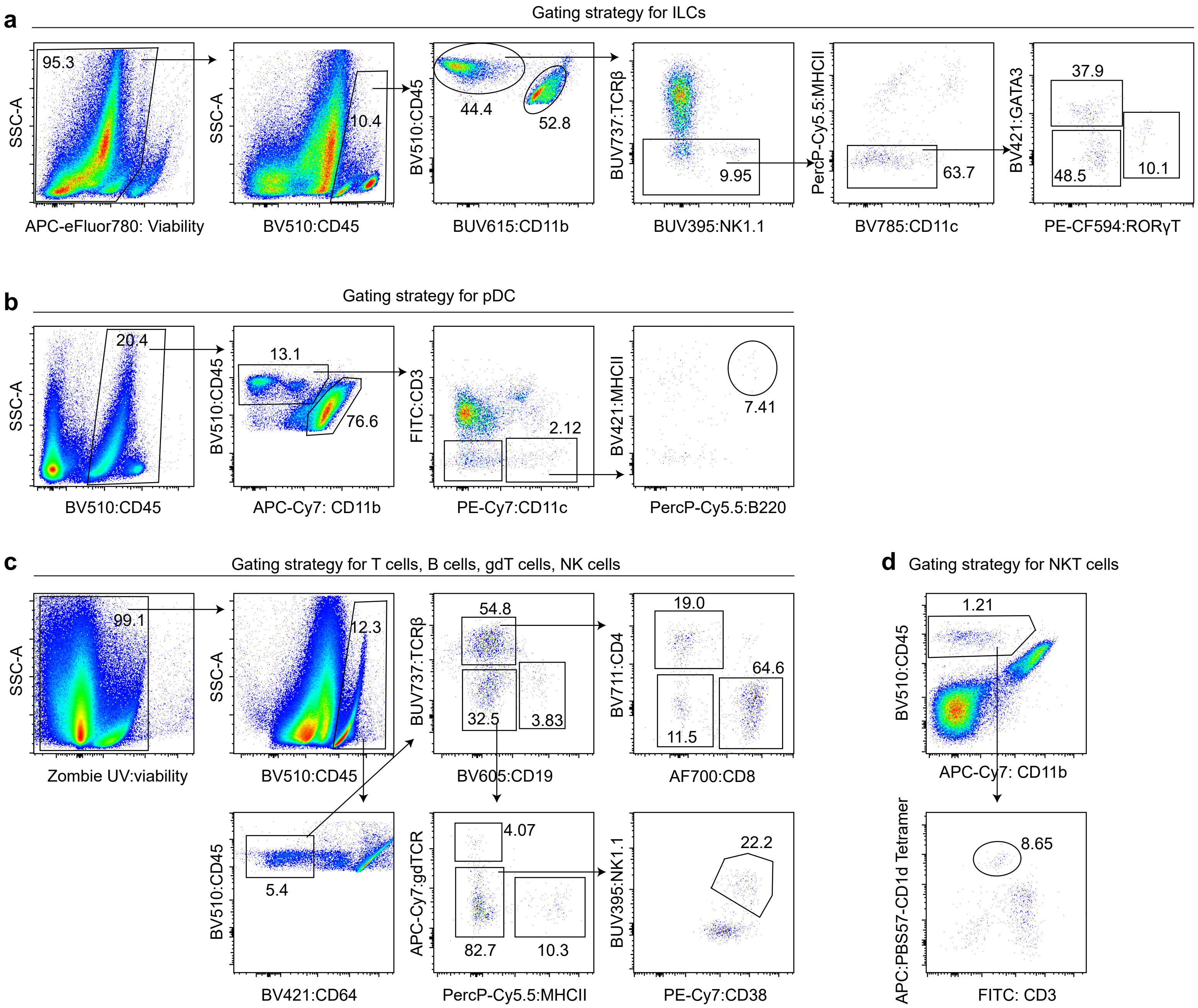

### Supplemental Fig8

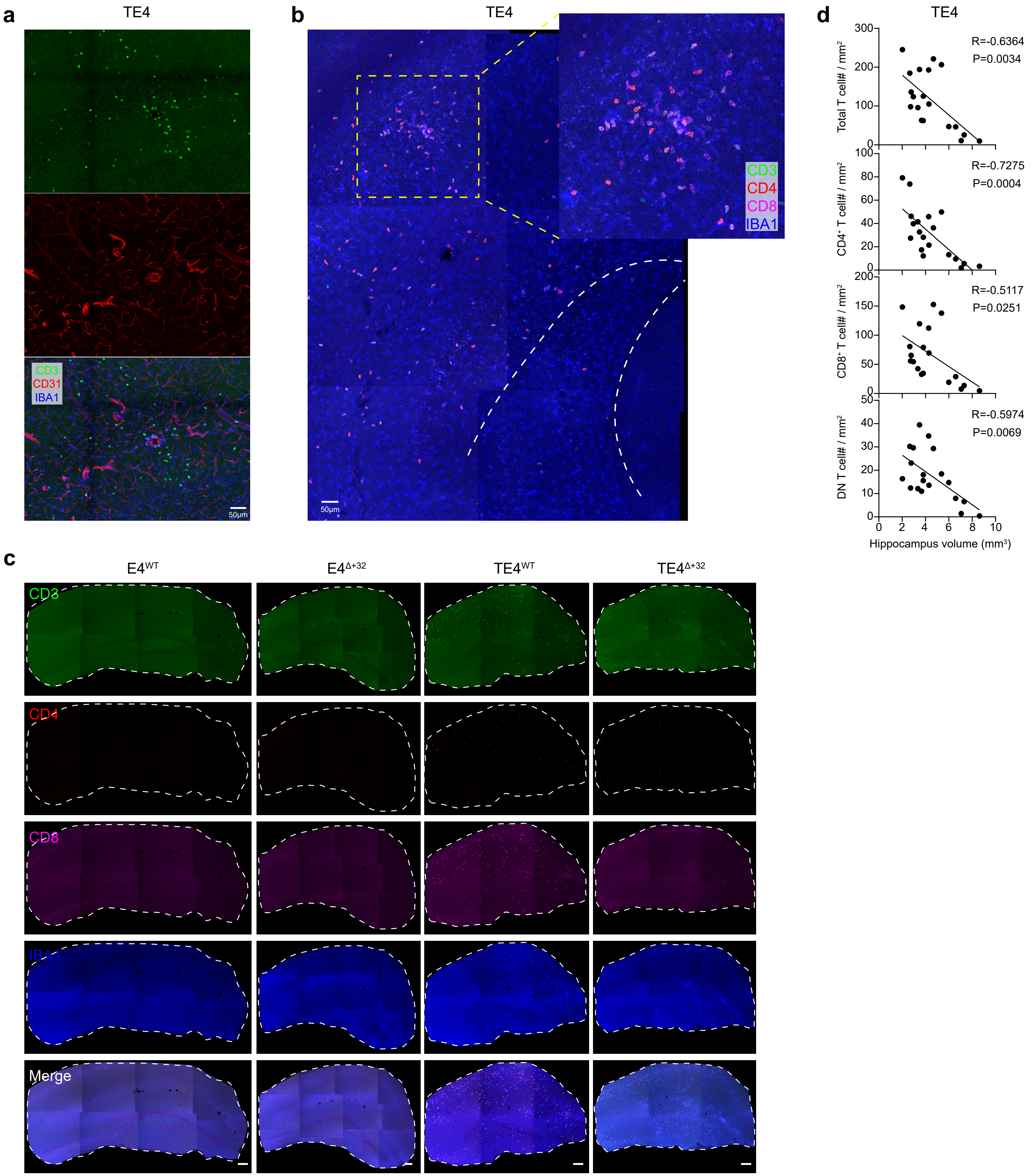

### Supplemental Fig9

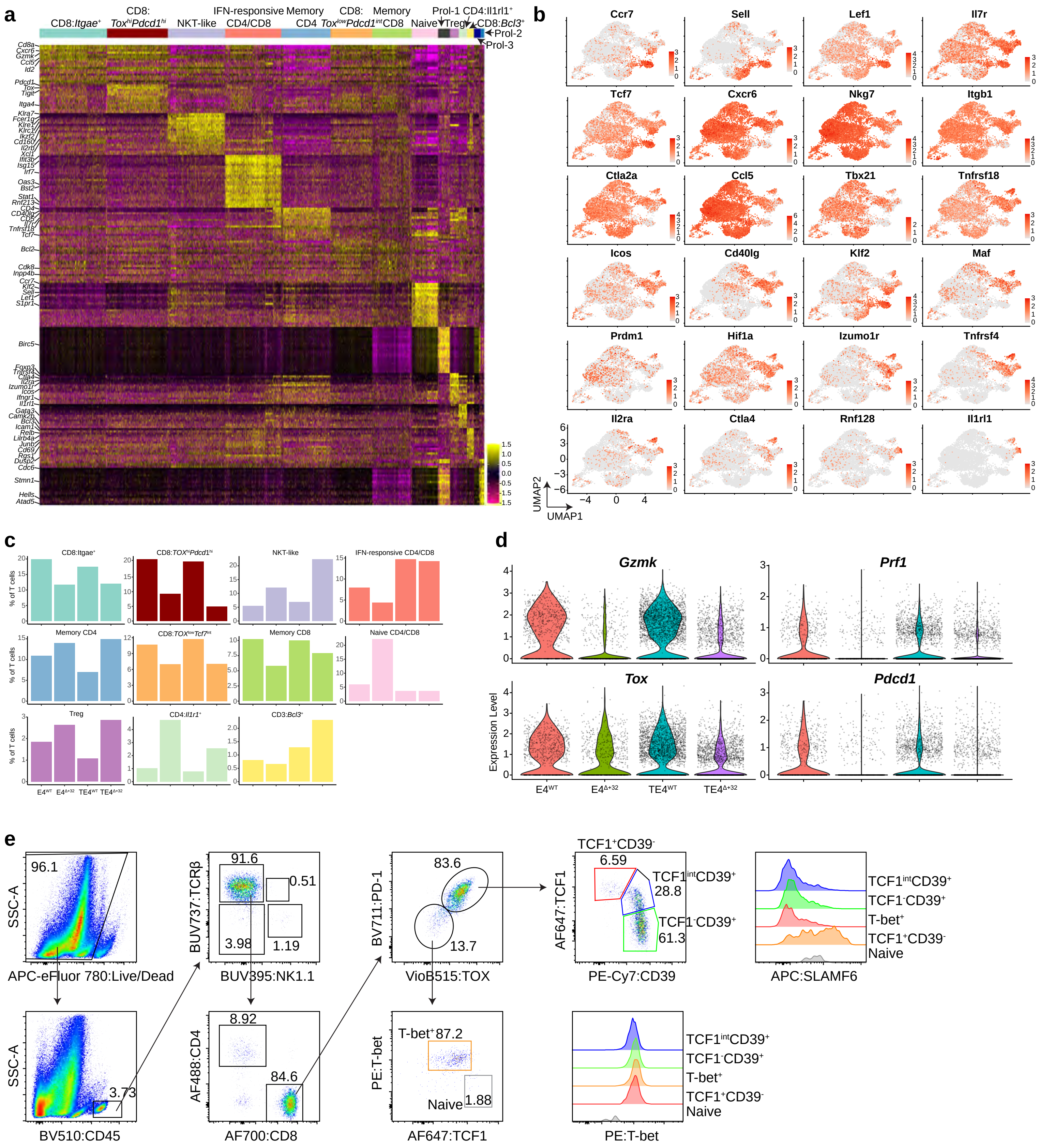

### Supplemental Fig10

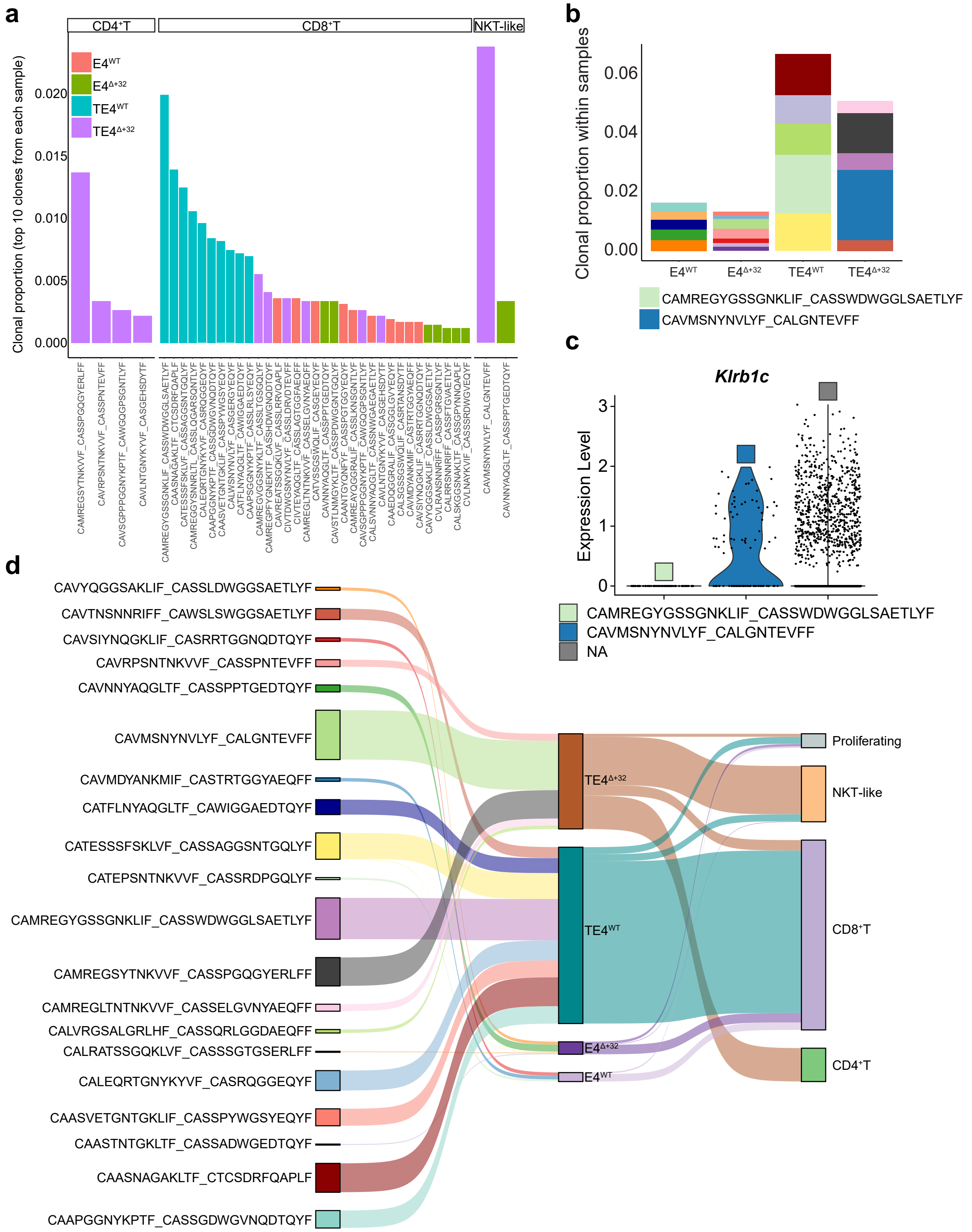

### Supplemental Fig11

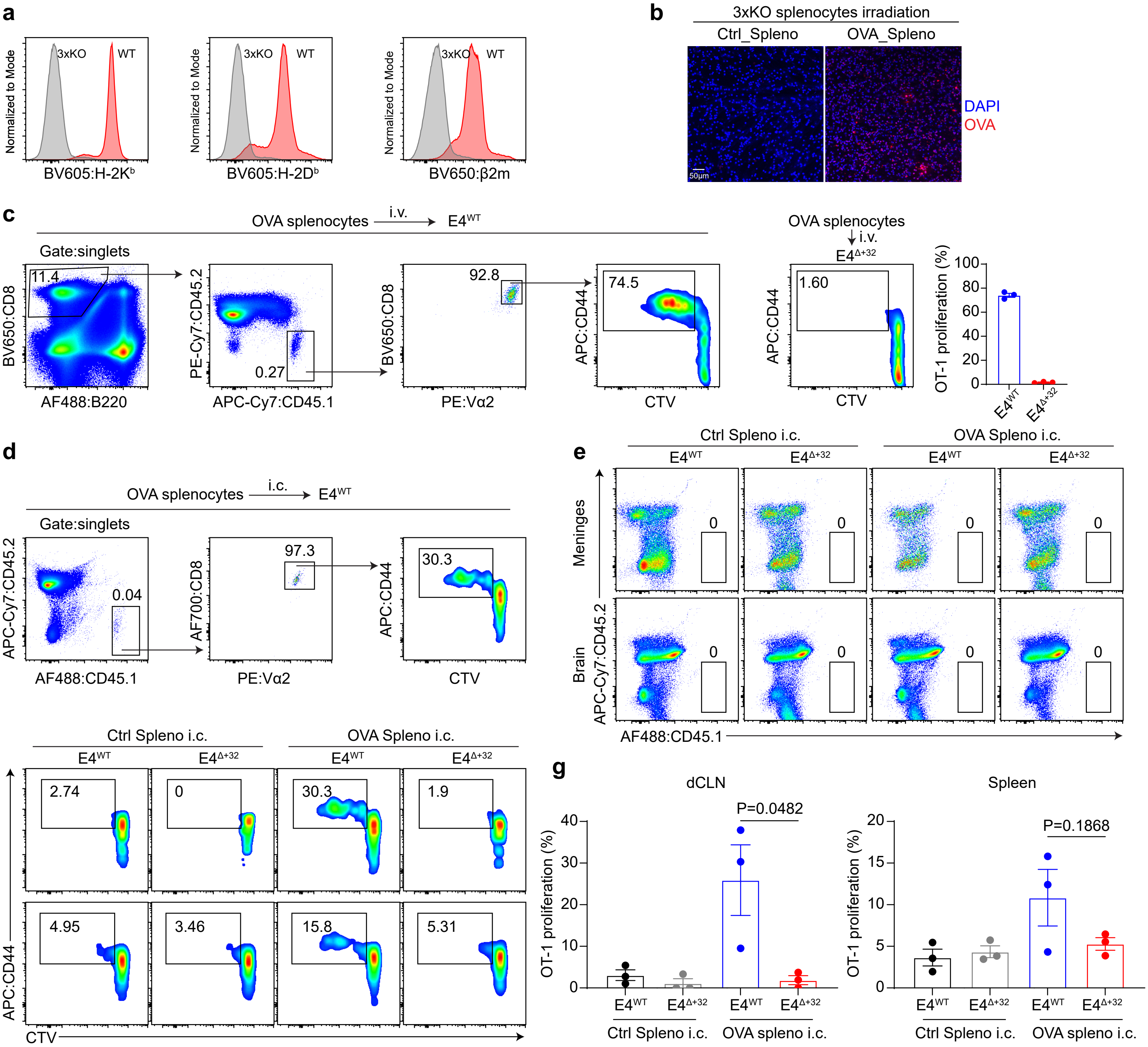
